# Enhancing the sensitivity of the envelope-following response for cochlear synaptopathy screening in humans: the role of stimulus envelope

**DOI:** 10.1101/2020.06.05.136184

**Authors:** Viacheslav Vasilkov, Markus Garrett, Manfred Mauermann, Sarah Verhulst

## Abstract

Auditory de-afferentation, a permanent reduction in the number of innerhair-cells and auditory-nerve synapses due to cochlear damage or synaptopathy, can reliably be quantified using temporal bone histology and immunostaining. However, there is an urgent need for non-invasive markers of synaptopathy to study its perceptual consequences in live humans and to develop effective therapeutic interventions. While animal studies have identified candidate auditory-evoked-potential (AEP) markers for synaptopathy, their interpretation in humans has suffered from translational issues related to neural generator differences, unknown hearing-damage histopathologies or lack of measurement sensitivity. To render AEP-based markers of synaptopathy more sensitive and differential to the synaptopathy aspect of sensorineural hearing loss, we followed a combined computational and experimental approach. Starting from the known characteristics of auditory-nerve physiology, we optimized the stimulus envelope to stimulate the available auditory-nerve population optimally and synchronously to generate strong envelope-following-responses (EFRs). We further used model simulations to explore which stimuli evoked a response that was sensitive to synaptopathy, while being maximally insensitive to possible co-existing outer-hair-cell pathologies. We compared the model-predicted trends to AEPs recorded in younger and older listeners (N=44, 24f) who had normal or impaired audiograms with suspected age-related synaptopathy in the older cohort. We conclude that optimal stimulation paradigms for EFR-based quantification of synaptopathy should have sharply rising envelope shapes, a minimal plateau duration of 1.7-2.1 ms for a 120-Hz modulation rate, and inter-peak intervals which contain near-zero amplitudes. From our recordings, the optimal EFR-evoking stimulus had a rectangular envelope shape with a 25% duty cycle and a 95% modulation depth. Older listeners with normal or impaired audiometric thresholds showed significantly reduced EFRs, which were consistent with how (age-induced) synaptopathy affected these responses in the model.

**Significance Statement:** Cochlear synaptopathy was in 2009 identified as a new form of sensorineural hearing loss (SNHL) that also affects primates and humans. However, clinical practice does not routinely screen for synaptopathy, and hence its consequences for degraded sound and speech perception remain unclear. Cochlear synaptopathy may thus remain undiagnosed and untreated in the aging population who often report self-reported hearing difficulties. To enable an EEG-based differential diagnosis of synaptopathy in humans, it is crucial to develop a recording method that evokes a robust response and emphasizes inter-individual differences. These differences should reflect the synaptopathy aspect of SNHL, while being insensitive to other aspects of SNHL (e.g. outer-hair-cell damage). This study uniquely combines computational modeling with experiments in normal and hearing-impaired listeners to design an EFR stimulation and recording paradigm that can be used for the diagnosis of synaptopathy in humans.

## Introduction

Noise overexposure, ototoxicity and aging can cause primary cochlear de-afferentation, i.e. progressive and irreversible damage to the afferent neuronal structures in the auditory periphery. One form of auditory de-afferentation is cochlear synaptopathy and refers to damaged synapses between the inner-hair-cells (IHCs) and auditory-nerve fibers (ANFs). This type of sensorineural hearing loss (SNHL) was first discovered in mouse models (Kujawa & Liberman, 2009) and has since been shown to affect macaques and humans as well (Wu et al., 2018; Viana et al., 2015; Valero et al., 2017). Cochlear synaptopathy specifically involves degeneration of the synaptic terminals of the spiral ganglion cells and precedes hair cell damage in the aging process (Wu et al., 2018; Fernandez et al., 2015; Sergeyenko et al., 2013). Furthermore, Ouabain and Kanic-acid treatment (Bourien et al., 2014; Shaheen et al., 2015; Chambers et al., 2016; Sheets, 2017) or noise-induced insults associated with temporary threshold shifts can cause cochlear synaptopathy (Kujawa & Liberman, 2009; Furman et al., 2013). The compelling histopathological evidence, along with outcomes from animal behavior studies of auditory de-afferentation (Schuknecht & Woellner, 1955; Lobarinas et al., 2013), have shown that cochlear synaptopathy has little effect on hearing sensitivity assessed through the behavioral audiogram or physiological threshold measures (e.g., distortion-product otoacoustic emissions; DPOAEs or auditory brainstem responses; ABRs). Cochlear synaptopathy might hence remain hidden during routine clinical hearing screening (Schaette & McAlpine, 2011), which typically assesses hearing sensitivity using the audiogram. We might hence overlook a large population of listeners who have accrued synaptopathy while their audiograms reflect normal hearing. Additionally, in listeners with age-related audiometric declines (ISO-7029:2000, 1991), only the hair-cell-damage aspect of SNHL is presently diagnosed and treated, while disregarding the possible co-existing synaptopathy aspect. To study the prevalence of synaptopathy, and its consequences for sound perception in humans, it is hence crucial to develop a non-invasive differential diagnostic test for synaptopathy as a first and necessary step towards effective therapeutic interventions.

The search for candidate non-invasive markers of synaptopathy has been ongoing since its discovery and has shown a promising role for auditory-evoked potentials (AEPs). Specifically, a reduction of the supra-threshold auditory brainstem response (ABR) amplitude was directly associated with histologically-verified cochlear synaptopathy (Kujawa & Liberman, 2009; Furman et al., 2013; Sergeyenko et al., 2013; Bourien et al., 2014; Möhrle et al., 2016a). Particularly, the ABR wave-I amplitude is currently considered as the most direct metric of cochlear synaptopathy (Kujawa & Liberman, 2009; Schaette & McAlpine, 2011; Lin et al., 2011; Furman et al., 2013; Prendergast et al., 2017a, 2018; Plack et al., 2016; Bramhall et al., 2019). The second measure proposed from animal cochlear synaptopathy studies is the envelope-following response (EFR), an AEP-type which is of predominant subcortical origin when the amplitude-modulation (AM) rate of the stimulus is above 80 Hz (Purcell et al., 2004). EFRs offer a more robust metric of cochlear synaptopathy than ABRs, as synaptopathy-induced EFR changes are greater than ABR amplitude reductions in the same animal (Shaheen et al., 2015; Parthasarathy et al., 2018).

Despite the compelling evidence from animal studies that combined AEP recordings with direct post-mortem synapse counts to diagnose synaptopathy, a direct translation of the identified markers toward a differential diagnosis in humans has proven difficult (Plack et al., 2016; Guest et al., 2017, 2018; Prendergast et al., 2017a, 2018; Bramhall et al., 2019; Garrett & Verhulst, 2019; Bharadwaj et al., 2019). The human data is not unambiguous in demonstrating reduced AEP metrics in listener groups with suspected synaptopathy (e.g., as induced through accumulated noise-exposure or age), and a number of studies report subtle or non-significant correlations between different electrophysiological markers that are sensitive to synaptopathy in animals (e.g. ABR and EFR amplitudes or slope changes, middle-ear-muscle reflex strength; Prendergast et al., 2017a; Guest et al., 2019; Garrett & Verhulst, 2019). Also individual performance differences in psychoacoustic tasks thought to be sensitive to cochlear synaptopathy (e.g. speech perception in noise, frequency discrimination, amplitude-modulation detection) do in some studies correlate with physiological markers of synaptopathy (e.g.; Bharadwaj et al., 2015; Mehraei et al., 2016; Liberman et al., 2016; Verhulst et al., 2018b), whereas in others they do not (e.g.; Schoof & Rosen, 2016; Guest et al., 2018; Prendergast et al., 2017b; Plack et al., 2014; Johannesen et al., 2019).

There are several aspects that contribute to these translational issues: the adopted physiological markers may be affected by species-specific biophysical processes (e.g. humans may be less vulnerable to noise damage than other species; Dobie & Humes, 2017; Valero et al., 2017; Hickox et al., 2017). Secondly, the markers may be differently impacted by different SNHL aspects, which may complicate their interpretation in terms of synaptopathy (Bramhall et al., 2019; Garrett & Verhulst, 2019). For example, OHC damage may result in a loss of compression which affects the level-dependent behavior of the markers, while a loss of high-threshold ANF fibers may reduce the dynamic range of stimulus intensities that can be represented. A third aspect relates to the limited extent by which cochlear synaptopathy might affect the considered the perceptual tasks (e.g. 50% ANF loss might be required to see a perceptual effect; Oxenham, 2016), and different ANF types may contribute differently to the considered electrophysiological markers. For example, low-spontaneous rate ANFs do not contribute strongly to the transient ABR wave-I (Bourien et al., 2014), but may contribute strongly to the EFR recorded to low-modulation depth stimuli (Bharadwaj et al., 2014). Lastly, it is possible that electrophysiological markers of synaptopathy simply have limited test-retest reliability for human use (D’haenens et al., 2008; Prendergast et al., 2018). However, before resorting to non-AEP based diagnostic markers, it is worthwhile to optimize existing AEP stimulation paradigms and analysis approaches to enhance the signal-to-noise ratio of the AEP, and consequently improve its test reliability. This route may yield a robust and sensitive diagnostic marker for auditory de-afferentation in humans, and help resolve the role of cochlear synaptopathy in future sound perception studies.

To address the above translational issues, this study aims to optimize stimulation and analysis paradigms to yield a reliable EFR-based cochlear synaptopathy diagnosis in humans. The EFR is an AEP-type evoked by repeated tokens of speech, noise or tonal stimuli and its strength reflects how well the auditory system represents the stimulus envelope. We particularly focus on auditory steady-state responses (ASSR) evoked by sustained periodic stimuli with constant carrier and modulation frequencies. In a clinical context, ASSRs are often used to assess hearing sensitivity whereas this study focuses on supra-threshold hearing. For this reason, we will adopt the EFR nomenclature which is common in cochlear synaptopathy studies.

This study draws from functional IHC-AN and peripheral auditory processing properties to develop stimulation paradigms that better reflect the available ANF population. This is important because a strong baseline EFR response will be more sensitive to changes induced by alterations in the ANF population. At the same time, we will adopt an optimized analysis method that extracts all the relevant envelope-following components from the raw EEG recordings. Furthermore, we explore how OHC functionality affects EFR generators to different stimulation paradigms. This will enable us to evaluate which EFR markers are maximally sensitive to synaptopathy, even when OHC damage is simultaneously present. We will test our biophysically-inspired stimulation paradigms in a computational model of the human auditory periphery (Verhulst et al., 2018a; Osses Vecchi & Verhulst, 2019, v1.2) that simulates EFRs for several frequency-specific SNHL profiles with different combinations of synaptopathy and OHC damage. The model simulations are compared against reference data recorded from three groups of study participants: young and older listeners with normal or elevated audiometric thresholds. The latter two groups show age-related SNHL (possibly including synaptopathy) with less or more OHC damage. We use the outcomes from the modeling and experimental study to identify which stimulus modifications and analysis paradigms increase the EFR strength and yield a sensitive non-invasive marker of cochlear synaptopathy for use in humans.

## Materials and Methods

### Stimuli

All AEP stimuli were generated in MATLAB R2015b (The MathWorks Inc., 2015) and had a sampling rate of 48 kHz for the recordings and 100 kHz for the model simulations. We designed and selected our stimuli on the basis of known observations in psychoacoustic and physiological studies of AM (e.g., van de Par & Kohlrausch, 1997; John et al., 2002, 2003; Bernstein & Trahiotis, 2002, 2009; Stürzebecher et al., 2003; Griffin et al., 2005; Dreyer & Delgutte, 2006; Laback et al., 2011; Klein-Hennig et al., 2011; Greenberg et al., 2017; Van Canneyt et al., 2019), and hypothesize that overall stronger EFRs might render individual EFR differences more robust for diagnostic purposes. From the results presented in the named studies, we suspect that a combination of increased silence gaps between the stimulus peaks and shorter stimulus duty cycles might cause more synchronized ANF activity, which, in turn might result in stronger EFRs. To test this hypothesis, we designed seven stimulus conditions with the same AM rate (120 Hz) and stimulus duration (400 ms), but with different carrier types (tone or noise), stimulus levels or envelope shapes. To validate our predictions and study the relative contribution of synaptopathy and OHC deficits, we simulated singleunit ANF responses as well as EFRs, the latter were also recorded in 44 participants. We used the widely-adopted sinusoidal amplitude-modulation (SAM) stimulus as the reference condition:

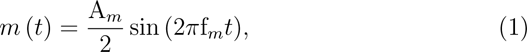

where A*_m_* corresponds to the peak-to-peak amplitude, f*_m_* is the modulation frequency and *t* is the time vector. Two carrier types were considered: a 4-kHz pure tone (PT) and a white-noise carrier with a 50-16000 Hz bandwidth (BB). Amplitude modulation was implemented by multiplying the carrier *c*(*t*) with [1 + md *∗ m*(*t*)*/*(A*_m_*)], where md = A*_m_/*A*_c_* and A*_c_* is the peak-to-peak amplitude of *c*(*t*). Figure 1a represents two cycles of the reference SAM stimulus with the 4-kHz PT carrier presented in sine phase. We additionally designed five AM stimuli with the same 4-kHz pure tone carrier, but with different modulators or sound levels.

**Figure 1.**
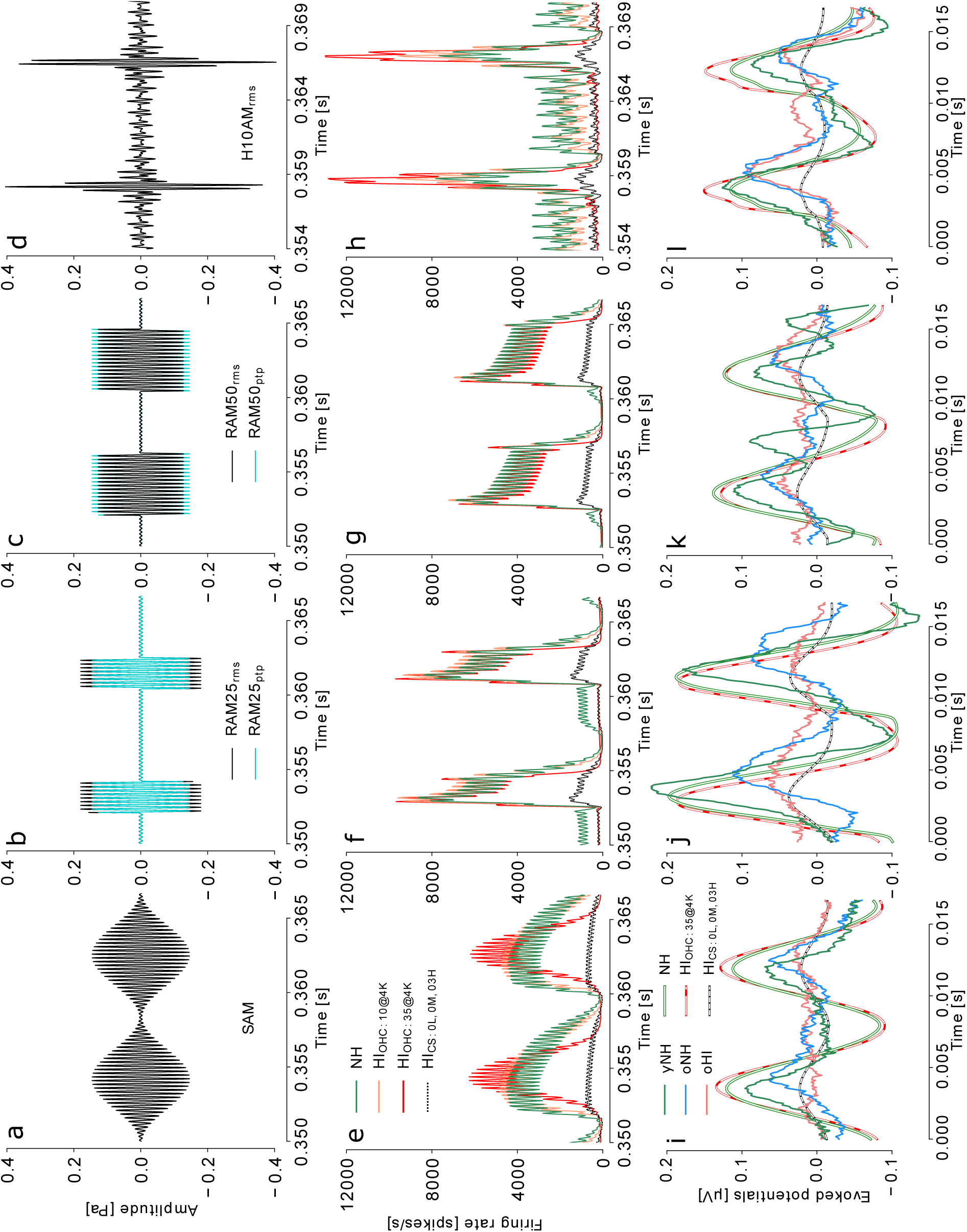
**a-d** Two cycles of the amplitude-modulated stimuli with different envelope shapes but the same modulation rate of 120 Hz. All stimuli were presented with the same RMS SPL (black) and stimuli with rectangular envelopes were additionally presented in an equal ptp amplitude to the reference SAM tone (cyan). **e-h** Simulated ANF responses at the 4-kHz CF evoked by the corresponding stimuli (equal-RMS). Solid traces depict responses summed across 19 AN fibers per IHC (i.e. intact ANF profile: 3 L, 3 M, 13 HSR fibers) and dotted black lines represent summed responses from three fibers per IHC (i.e. severe synaptopathy HI_CS_: 0 L, 0 M, 3 HSR fibers). HI_OHC:10@4K_ and HI_OHC:35@4K_ traces represent responses for simulated sloping audiometric hearing loss with 10 dB or 35 dB threshold elevation at 4 kHz. NH shows the responses without simulated hearing deficits. **i-l** Time-domain representation of simulated and recorded EFRs in response to stimuli with different envelope shapes (same RMS). Open traces depict simulated EFRs for normal-hearing (NH), extreme synaptopathy (HI_CS:0L,0M,3H_) and audiometric (HI_OHC:35@4K_) profiles. Solid traces depict recorded EFRs averaged across young (yNH) and old (oNH) normal-hearing, and old hearing-impaired (oHI) participants groups.

The second, non-sinusoidal, periodic modulator we considered, had a rectangular waveform which was presented with a period of 2*π* and a 25% duty cycle (RAM25, Fig. 1b). The RAM25 modulator was generated using the *square(t, 100d)* function and can be described using the Fourier series expansion:

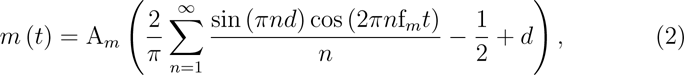

where d = 0.25 denotes the duty cycle, i.e. the ratio between the pulse width and the total period of the modulator, n is the harmonic number of the series.

The third modulator was a rectangular waveform with a 50% duty cycle (RAM50; Fig. 1c) with d=0.5:

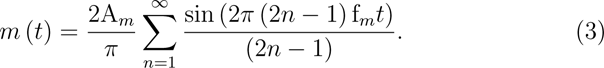

The fourth modulator was a ten-harmonic complex (H10AM, Fig. 1d) presented in cosine phase:

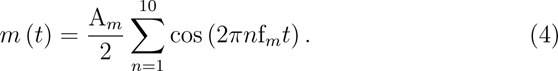

The different AM stimuli were calibrated to have the same root-mean-square (RMS) sound pressure of 70 dB SPL. To study whether there was an effect of RMS versus peak-to-peak sound calibration, two stimuli were also calibrated to have the same peak-to-peak amplitude as the reference SAM tone (i.e. RAM25*_ptp_* and RAM50*_ptp_*; Fig. 1b and Fig. 1c, cyan). They were presented at 68.18 dB SPL and 71.18 dB SPL, respectively. All stimuli were 95% amplitude modulated (e.g., −0.45 dB re 100% modulation) with a starting phase shift of 3*π*/2. An exception was the H10 complex modulator, which had a 0 starting phase. Each stimulus had gradual on and offsets (2.5% tapered-cosine Tukey window) and was presented 1000 times using 500 repetitions per polarity. Stimuli were presented monaurally and a uniformly distributed inter-stimulus silence interval of 100 *±* 10 ms was applied.

**ABR**s were recorded to 3000 repetitions of a 80-*µ*s click presented monaurally with alternating polarity at a mean rate of 10 Hz (including the uni-formly distributed 10% silence jitter). Three stimulus levels were tested (70, 85, and 100 dB peSPL) and we only considered the 70 and 100 dB peSPL conditions conditions for this study. ABR and EFR stimuli were calibrated using an oscilloscope, ear simulator (Brüel & Kjær 4157) and sound level meter (Brüel & Kjær 2610).

### Model of the auditory periphery

The auditory periphery model we adopted (Verhulst et al., 2018a; Osses Vecchi & Verhulst, 2019, model implementation v1.2) simulates auditory processing along the ascending pathways (Fig. 3) and includes middle-ear filtering, a nonlinear transmission-line representation of human cochlear mechanics (Verhulst et al., 2012; Altoé et al., 2014), a model of the IHC-AN complex (Altoé et al., 2018), and a phenomenological description of ventral cochlear nucleus (CN) and inferior colliculus (IC) neurons (Nelson & Carney, 2004). The model reasonably captures properties of AN fiber types with different spontaneous rates, level-dependent ABR/EFR characteristics, and furthermore can simulate frequency-specific hearing impairments related to OHC damage and synaptopathy (Verhulst et al., 2015, 2016, 2018a).

Cochlear synaptopathy was modeled by reducing the number of IHC-AN synapses of different ANF types at each simulated tonotopic location. The normal-hearing (NH) model had 19 fibers with three spontaneous-rate (SR) fibers synapsing onto each IHC (Verhulst et al., 2018a): 3 low (L), 3 medium (M) and 13 high (H) SR fibers, following the ratio observed in cats (Liber-man, 1978). Three synaptopathy profiles were implemented by removing the following fiber types across the tonotopic axis: (i) all LSR and MSR fibers (HI_CS:0L,0M,13H_), (ii) all LSR, MSR and 50 % of the HSR fibers (HI_CS:0L,0M,07H_), and (iii) all LSR, MSR and 80 % of the HSR fibers (HI_CS:0L,0M,03H_). We limited our simulations to uniform, CF-independent synaptopathy profiles. No IHC-specific dysfunctions were simulated in the current study as synaptopathy was suggested to occur without destroying the sensory cells (Kujawa & Liberman, 2009; Lin et al., 2011; Furman et al., 2013; Shaheen et al., 2015). However, IHC loss can be simulated by introducing a complete synaptopathy (0L, 0M, 0H SR fibers).

OHC dysfunction caused by damaged mechano-receptors or presbycusis was simulated by adjusting the gain parameters of the cochlear model filters to yield frequency-specific cochlear gain loss profiles. Figure 2a shows mean audiometric thresholds of the study participants along with corresponding simulated cochlear gain loss profiles (dashed and solid lines, respectively). These gain loss profiles (in dB HL) were used to determine the parameters of the cochlear filter gain relative to the normal-hearing cochlear filter gain at CFs corresponding to the audiometric testing frequencies (see Fig.2 in Verhulst et al., 2016). Even though the model can simulate individual cochlear gain loss profiles in great detail, we limited our simulations to a range of sloping high-frequency profiles approximating the average audiograms of each participant group (yNH, oNH, oHI; Fig. 2a).

**Figure 2.**
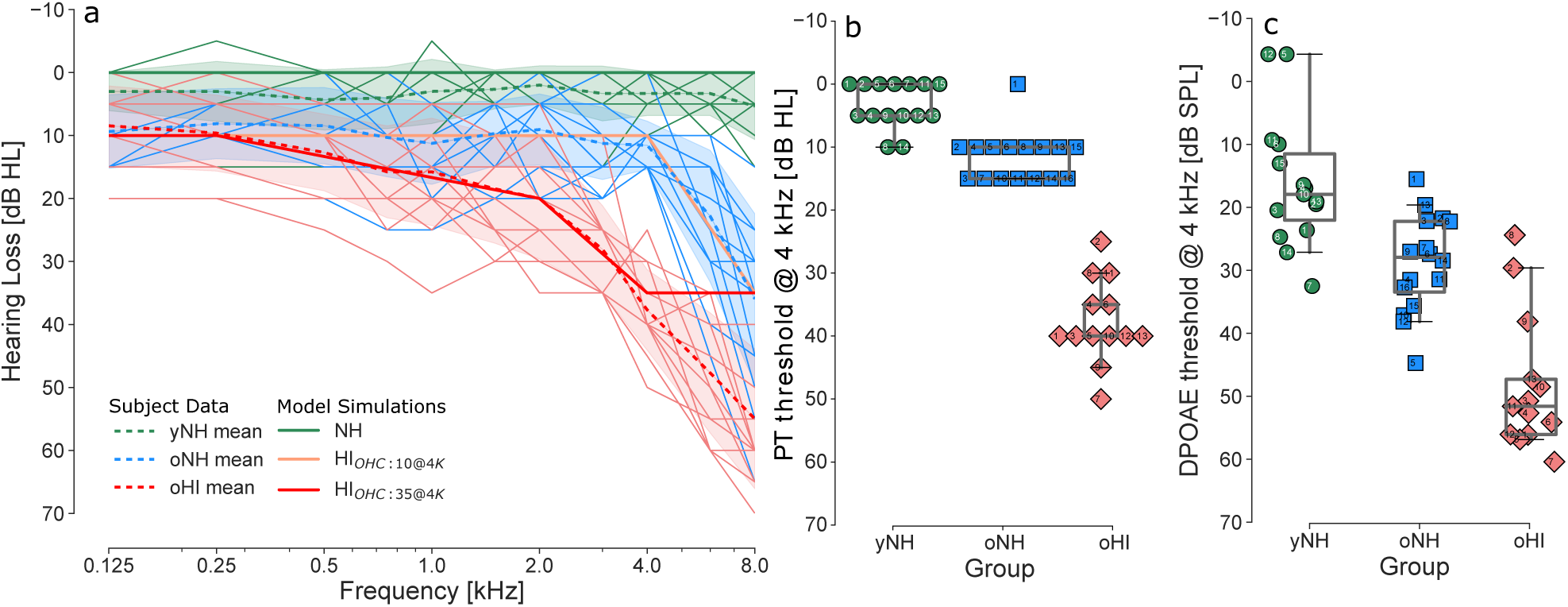
**a** Pure-tone hearing thresholds measured at frequencies between 0.125 and 8 kHz. Dashed traces depict mean values across yNH, oNH and oHI groups. Solid lines represent simulated cochlear gain loss profiles corresponding to the mean experimental audiograms (NH, HI_OHC:10@4K_ and HI_OHC:35@4K_). **b** Pure-tone hearing thresholds and **c** distortion-product otoacoustic emission thresholds at 4 kHz.

AEPs were simulated by adding up instantaneous firing rates across a tono-topic array of 401 IHC-AN/CN/IC units (Verhulst et al., 2018a) positioned along the cochlea according to the frequency-position map (Greenwood, 1990). Responses from 19 AN fibers of three SR types which synapse onto a single IHC were summed at each CF to form the input to a single CN unit of the same CF iin the NH model. The instantaneous firing rate of a single CN unit served as input to a single IC unit. A same-frequency inhibition and excitation model for the CN and IC units was adopted and captures the modulation filtering and onset enhancement characteristics of auditory brainstem and midbrain neurons (Nelson & Carney, 2004). Instantaneous firing rates were summed across all simulated CFs for three processing stages: (i) the AN (after summing 19 ANF responses for each IHC across the different CFs) to yield the W-I response; (ii) the CN and (iii) IC model stages yielding the W-III and W-V response respectively (Fig. 3). For EFR simulations, the population responses from the AN, CN and IC processing stages were added to realistically capture the different subcortical sources that contribute to EFRs (Dolphin & Mountain, 1992; Kuwada et al., 2002).

**Figure 3.**
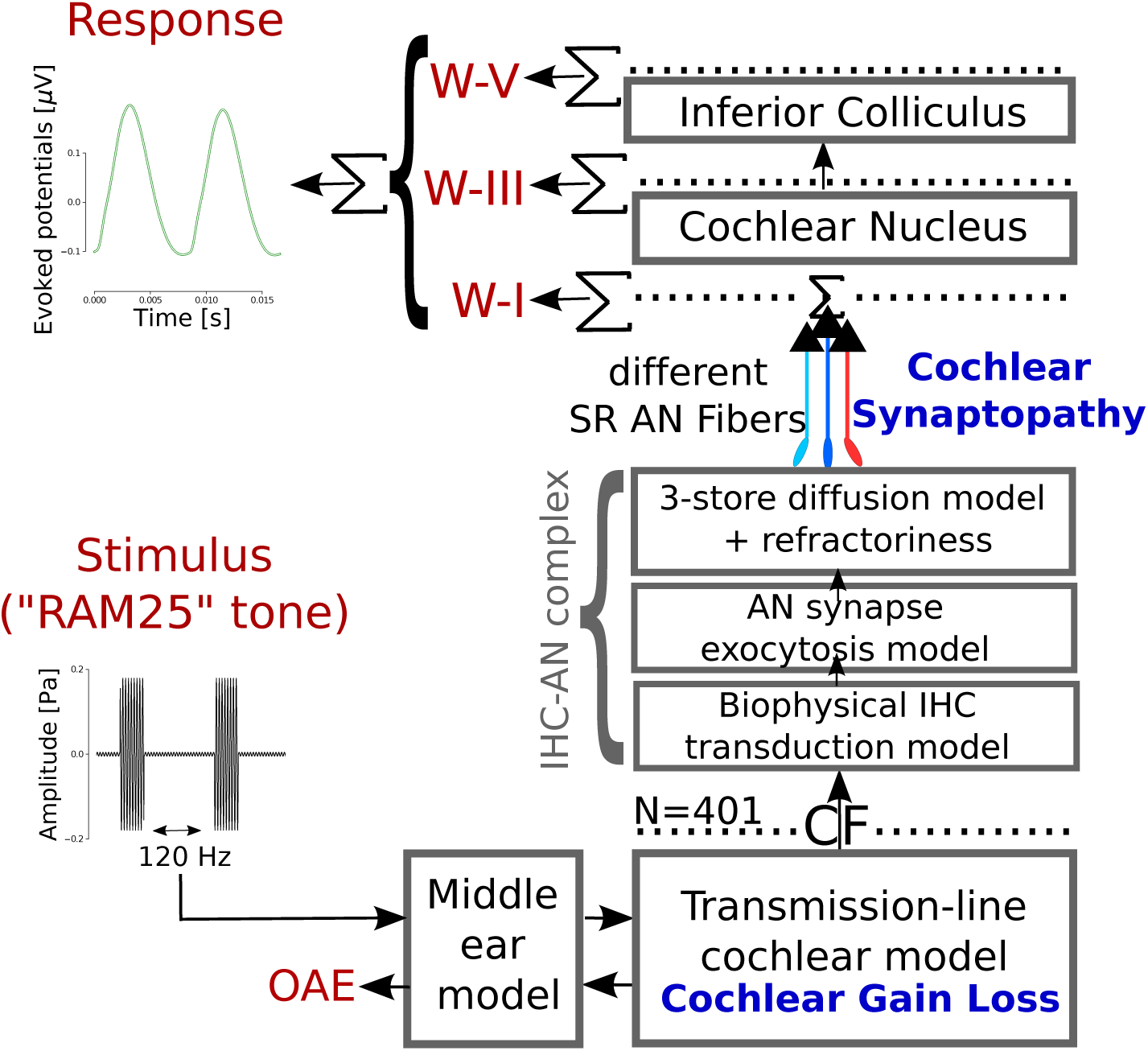
Schematic of the adopted computational model of the auditory periphery that simulates subcortical sources of human AEPs in response to acoustic stimuli (Verhulst et al., 2018a; Osses Vecchi & Verhulst, 2019).

### Participants

A total of 44 participants were recruited into three groups based on the combination of two criteria: age and audiometric profile. The audiometric thresholds, sex and ages of all participants are listed in Table A1 and Figure 2 depicts pure-tone thresholds for the audiometrically-better ear (which was used for the experiments). Participants were informed about the ex-perimental procedures and the experiments were approved by the ethical commission of the University of Oldenburg. They gave a written informed consent and were paid for their participation.

The young normal-hearing (yNH) group consisted of 15 participants (8 females; age *±* standard deviation: 24.5 *±* 2.2), who had pure-tone hearing thresholds below 25 dB HL across the standard audiometric frequency range. The older normal-hearing (oNH) group comprised 16 participants (8 females) with ages between 60-70 years (64.3 *±* 1.8 y/o), and had normal-hearing thresholds below 25 dB HL across the 0.125-4 kHz frequencies. The older hearing-impaired (oHI) group consisted of 13 participants (8 females) with ages between 60-70 years (65.2 *±* 1.8 y/o) and had sloping high-frequency audiograms that exceeded 25 dB HL at 4 kHz. An otoscopic inspection was performed to ensure that participants had no visible pathologies or obstructions. Audiograms were measured for standard frequencies between 0.125-8 kHz using a clinical audiometer (Auritec AT 900) and over-ear audiometric headphones (Sennheiser HDA 200). Figure 2b shows individual hearing thresholds at 4-kHz (which corresponds to the PT carrier frequency of the EFR stimuli) for yNH (3.3 *±* 3.5 dB HL), oNH (11.6 *±* 3.8 dB HL) and oHI (37.7 *±* 6.4 dB HL) groups.

Additionally, we recorded distortion-product otoacoustic emissions (DPOAEs) to quantify the OHC-related aspect of SNHL. To this end, ER-2 insert earphones were coupled to the ER-10B+ microphone system (Etymotic Re-search) and we used a custom-made MATLAB program (Mauermann, 2013) for DPOAE recording and analysis. Primary tone pairs were simultaneously presented with a fixed f_2_*/*f_1_ ratio of 1.2 using a continuously sweeping DPOAE paradigm (Long et al., 2008). Primary-frequencies were exponentially swept (2 s/octave) during stimulus presentation over a 1/3 octave range around the geometric mean. f_2_ ranged from 1 to 4 kHz using octave steps. Levels were set according to the “scissors” level paradigm (L_1_ = 0.4 *·* L_2_ +39 dB; Kummer et al., 1998), using a step size of 6 dB. L_2_ ranged between 30-60 dB SPL for yNH and oNH participants, and between 30-72 dB SPL for oHI participants. DPOAE thresholds were derived from recorded DPOAE-L_2_-level series for each mean f_2_ frequency within the measured frequency range using a bootstrapping procedure. In this procedure, a tailored cubic growth function was fit through the DPOAE-L_2_ data points to determine the DPOAE threshold (Verhulst et al., 2016) using a resampling method: 200 growth functions were calculated from random draws within each confidence interval of the DPOAE-L_2_ data points. Using this method, it is possible to calculate the standard deviation of the fits and DPOAE thresholds, given the standard deviations of the DPOAE data-points in the level series. DPOAE thresholds were determined as the median of the L_2_ levels at which the mean curve fit reached a level of −25 dB SPL (Boege & Janssen, 2002) for each bootstrap average. DPOAE thresholds at 4 kHz are depicted in Fig. 2c for yNH (16.1 *±* 10.0 dB), oNH (28.9 *±* 7.5 dB) and oHI (48.2 *±* 10.5 dB) groups.

### Recording setup and preprocessing

Measurements were performed in a double-walled electrically-shielded booth while participants sat comfortably in a reclining chair and watched a silent movie. Stimuli were presented monaurally (using the audiometrically better ear) over magnetically-shielded ER-2 insert earphones (Etymotic Research) connected to a TDT-HB7 headphone driver (Tucker-Davis) and a Fireface UCX sound card (RME). EEG data were recorded using a 64-channel recording system (BioSemi) and BioSemi Active-electrodes which were spaced equidistantly in an EEG recording cap (EasyCap). A common-mode-sense active electrode was placed on the fronto-central midline and a driven-right-leg passive electrode was placed on the tip of the nose of the participant. Reference electrodes were placed on each earlobe. A 24-bit AD conversion with sampling rate of 16384 Hz was used to digitize and store the raw data (for additional setup details see Garrett et al., 2019). The raw data were preprocessed using Python (version 2.7.10) and the MNE-Python (version 0.9.0) open-source software package (Gramfort et al., 2013, 2014). The vertex (Cz) channel potentials were re-referenced to the off-line-average of the two earlobe channel potentials to obtain the AEP. Pre-processed time-domain AEP waveforms were further processed in MATLAB R2014b (The MathWorks Inc., 2014) to perform waveform averaging, bootstrapping and feature extraction.

### ABR analysis

ABR recordings to positive and negative polarity clicks were high-pass filtered with a cut-off frequency of 200 Hz and then low-pass filtered with a cutoff frequency of 2000 Hz using a zero-phase filter (4th order IIR Butterworth filter). ABR recordings were epoched into 20 ms windows relative to the stimulus onset. Bad epochs were identified using the joint probability criteria as implemented in EEGLAB (Brunner et al., 2013).

ABR waveforms and variability were estimated using a bootstrap procedure. For each condition, 1000 time-domain epochs (for positive and negative stimulus polarities) were randomly drawn with replacement and averaged 200 times. ABR wave-I and wave-V waveform peaks were identified through visual inspection. ABR amplitudes [in *µ*V] were defined as the absolute amplitude difference between a positive peak and corresponding maximal negative deflection before the next up-going slope (Picton, 2010).

### EFR analysis

EFR recordings were epoched to 400-ms windows starting from the stimulus onset and were baseline corrected using the average amplitude per epoch. EFR magnitudes were derived by estimating the amplitude of a time-domain response which predominantly contained stimulus-driven energy. This signal was obtained by removing the individual electrophysiological noise floor (NF) and stimulus-irrelevant EEG components (Fig. 4). We calculated the mean EFR magnitude and corresponding standard deviation across the available epochs using a bootstrap procedure (Zhu et al., 2013). In each bootstrap run, we calculated a magnitude spectrum (in *µ*V) using the FFT of the time-domain average of 1000 randomly sampled (with replacement) response epochs, i.e. 500 epochs per stimulus polarity. Epochs were windowed using a 2% tapered-cosine (Tukey) window before applying the frequency-domain transformation. An example of an EEG magnitude spectrum for one boot-strap average and corresponding NF estimates is shown for a listener from the NH group in Fig. 4a. To include all energy related to the stimulus envelope, the EFR magnitude was computed from the spectrum based on the energy at the frequency corresponding to the stimulus modulation rate (f_1_=120 Hz) and its harmonics (f_(_*_k_*_)_=*k*\*f_1_, *k*=[1..5] for our recordings) using the energy above the NF. The noise floor of f_1_-f_5_ was computed as the average magnitude across the ten bins centered around the corresponding frequency (5 bins on either side). Spectral peaks at f_1_-f_5_ (F*_n_*) were then corrected by subtracting the respective NF*_n_* values to yield a relative peak-to-noise-floor (PtN) magnitude estimate (blue arrows; Fig. 4a). Negative PtN estimates (i.e., when spectral peaks F*_n_* were smaller than the noise-floor NF*_n_*) were set to zero and energy at other frequencies was removed before constructing the EFR waveform. To yield a time-domain EFR signal which mostly contains energy related to the stimulation AM, we performed an iFFT using the noise-floor corrected spectral peaks (F*_n_*-NF*_n_*) and their corresponding phase angles (*θ_n_*). This procedure focuses on the individual NF-corrected component of the recording and uses absolute signal values (in *µ*V) rather than SNR values which can be affected by NF level variability between NH and HI listeners. Figure 4b depicts the comparison between the recording and reconstructed EFR waveform (thin gray and solid blue traces, respectively). Note that the recordings in Fig. 4b were band-pass filtered between 117 Hz and 603 Hz to emphasize stimulus-envelope-related components and remove irrelevant energy beyond the modulation frequency and its harmonics for visual clarity. Finally, the EFR magnitude was defined as half the peak-to-peak amplitude of the reconstructed time domain-signal waveform, i.e:

**Figure 4.**
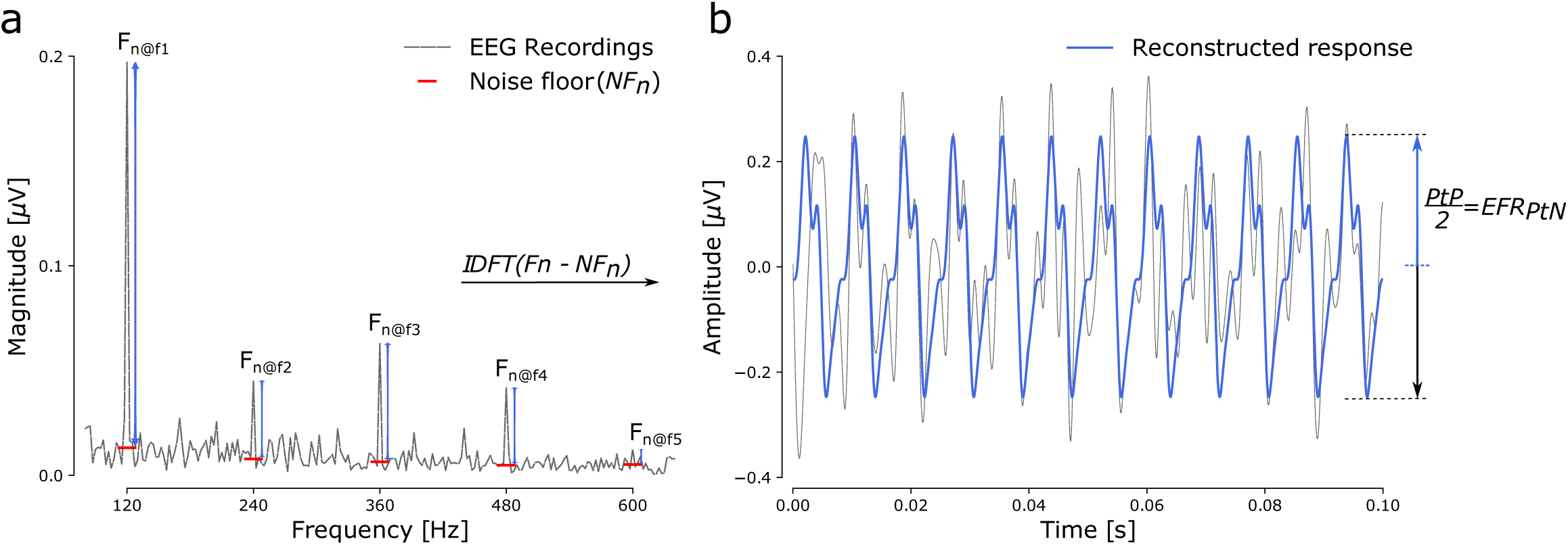
Illustration of how the EFR*_P tN_* was computed from the raw EEG recordings. Data correspond to yNH #7 **a** Magnitude spectrum (gray) of the AEP to a 70-dB-SPL RAM25*_rms_* stimulus. Red dash markers depict the estimated noise floor (NF) and blue vertical arrows indicate peak (F*_n_*) to NF*_n_* magnitudes at the modulation frequency and its harmonics. **b** 100 ms-scaled time-domain representation of the recorded AEP (gray) and reconstructed time-domain EFR waveform (blue) based on noise-corrected energy at the modulation frequency and harmonic components (F*_n_*-NF*_n_*). Time-domain reconstruction (iFFT) used the original F*_n_* phase angle and the single sided EEG magnitude spectrum. The EFR magnitude was defined as half the peak-to-peak amplitude of the reconstructed signal in the time domain (blue arrow) for each bootstrap run.

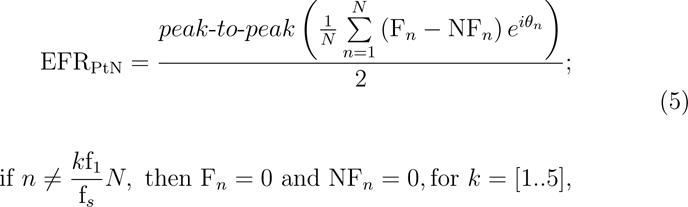

where *N* corresponds to the length of the magnitude spectrum, and f*_s_* the sampling rate. As a result of the bootstrapping procedure, we obtained 200 reconstructed time-domain waveforms for each listener and stimulus condition, which we used to accurately estimate the EFR_PtN_ magnitude and its standard deviation. Simulated EFR magnitudes for different SNHL profiles were directly derived from the time-domain responses, because no noise or stochastic processes were implemented in the adopted model version. Simulated EFR magnitudes were defined as half the peak-to-peak amplitude of the average one-modulation-cycle waveform across the 400-ms epoch duration.

## Results

### Simulated ANF responses to EFR stimuli with different envelope shapes

Figure 1e-h shows simulated ANF firing rates at the CF of 4 kHz, summed across the different fiber types. Responses to two cycles of the reference 70-dB-SPL SAM tone are shown, as well as responses to the stimuli with other envelope shapes. Each panel depicts simulations of the normal-hearing model (NH: green) and SNHL models: two degrees of OHC damage (HI_OHC:10@4K_: salmon, HI_OHC:35@4K_: red) and one synaptopathy profile (HI_CS:0L,0M,03H_: black). The NH simulations generally followed the stimulus envelope shape, but differences were seen in response strength and distortion. In particular, the NH SAM response (Fig. 1e) had a distorted shape with strong firing rates to the sloping parts of each stimulation cycle and short temporal regions without firing near stimulus envelope minima. Low firing rates occurred towards the end of each cycle due to the IHC-AN adaptation properties (Altoé et al., 2018).

Despite their similar modulation rates and SPL levels, the other stimuli evoked responses with steeper attack/decay slopes and broader temporal regions with near-zero firing rates (Fig. 1f,g). The RAM stimuli evoked stronger responses compared to the reference SAM condition and the re-sponse to the H10AM stimulus (Fig. 1h, green) was characterized by sharp peaks and pronounced firing to the small stimulus fluctuations between two stimulus cycle peaks. Comparison between the conditions shows that long inter-peak intervals (IPI) were a necessary condition to yield high peak firing rates during supra-threshold stimulation with stimuli of high modulation rates (Fig. 1f, green; RAM25). Longer IPIs may provide more time for the neuron to recover (e.g. replenish neurotransmitter) and allow it to respond more reliably to each stimulation cycle. Comparison between the RAM50 and RAM25 firing rates showed increased firing rates when the duty cycle of the stimulus envelope decreased and this is, in the model, caused by more synchronized spiking activity to the stimulation plateau and reduced spiking in the silence windows of the longer RAM25 IPI.

Cochlear amplification responded differently to the considered stimulus envelope shapes, and consequently, OHC damage was seen to influence the simulated ANF responses differently. Simulated NH ANF rates (green) and HI rates for 10 (blue) and 35 dB HL (red) loss at 4 kHz are depicted in Fig. 1e-h. For the reference SAM stimulus (Fig. 1e), increasing the degree of cochlear gain loss resulted in linearized and less distorted ANF firing rates and broader silence regions between the response cycles. Additionally, enhanced responses were observed near the stimulus envelope maxima in comparison to the NH responses. Stronger peak ANF rates for HI vs NH simulations were also observed for the H10AM condition and both were, in the model, caused by a combination of small ANF responses to stimulus minima and a cochlear compression loss, which together facilitated stronger onset ANF firing. A different pattern of ANF firing was observed for the RAM stimuli after simulating cochlear gain loss: ANF rates to all cochlear gain loss profiles largely overlapped and showed only minor differences between the peak rates. The RAM-ANF responses were thus only marginally impacted by simulated OHC damage, and in the model, this behavior resulted from the long, near-zero, IPIs followed by sharply rising envelope onsets, which together minimize the time the cochlear amplification has to impact the temporal response.

Introducing synaptopathy drastically reduced the firing rates to all stimuli (Fig. 1e-h, black dashed lines) and the reduction was proportional to the remaining number of intact ANFs (see also Fig 5). Figure 1e-h shows that the stimulus envelope shape has an important effect on how ANF firing patterns were affected by different aspects of SNHL. Responses to the SAM and H10AM stimuli showed that inter-peak envelopes with low sound intensities can yield stronger peak responses after OHC damage that, to a certain degree, can compensate for the reduced firing rates caused by synaptopathy when both SNHL aspects are present. In contrast, the ANF rates to the RAM stimuli were strongly affected by synaptopathy, but not OHC damage. These ANF response simulations at CF hence suggest that AM stimuli with rectangular envelope shapes can provide a differential and enhanced sensitivity to the synaptopathy aspect of SNHL.

**Figure 5.**
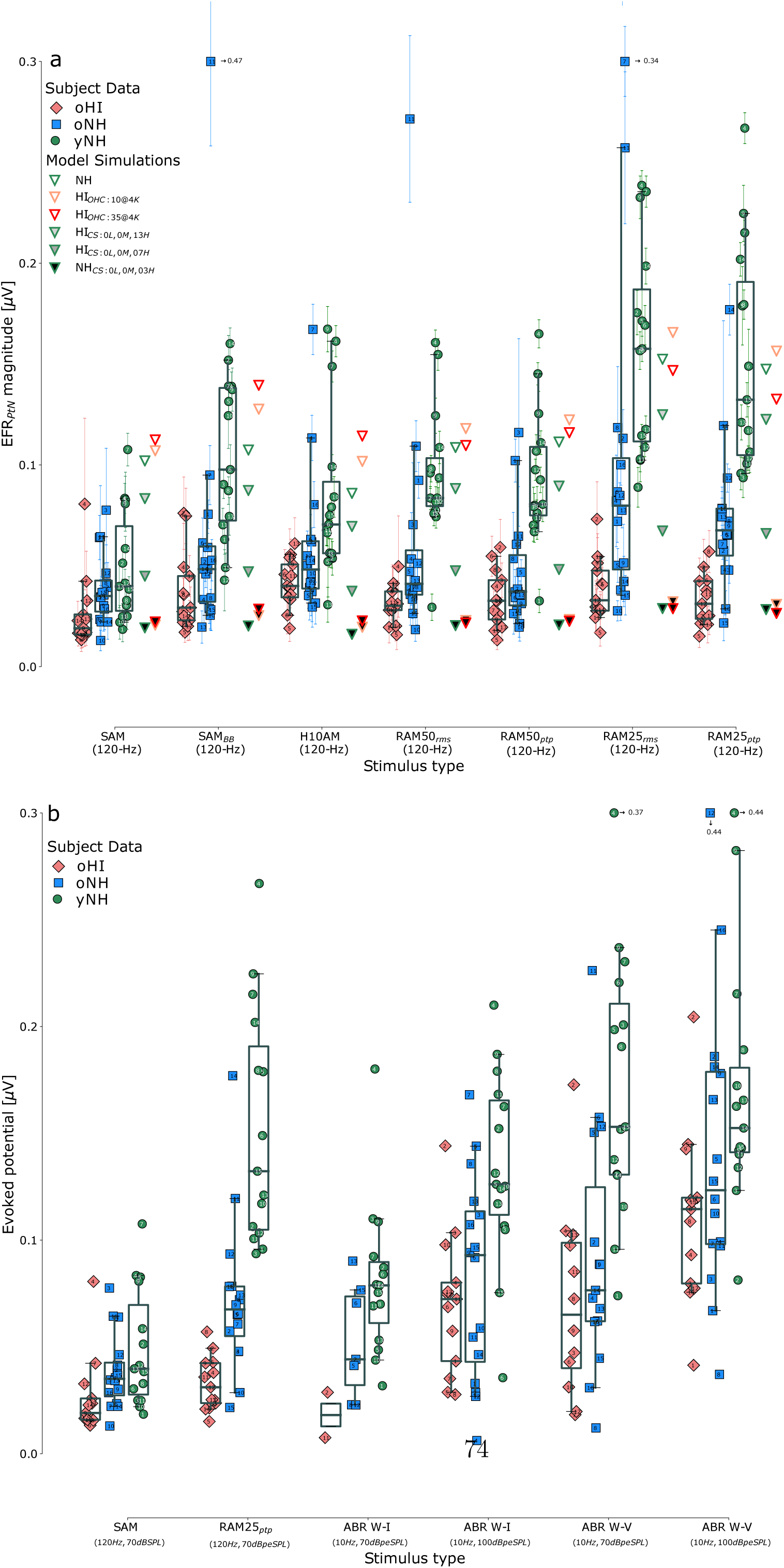
Recorded AEPs for individuals of the yNH (green circles), oNH (blue squares) and oHI (red diamonds) subgroups along with model simulations (triangles) for different SNHL profiles and stimulus types. **a** EFR*_P tN_* magnitudes evoked by amplitude-modulated stimuli with different envelope shapes. **b** Comparison between EFRs and transient ABR waveform features to 10-Hz click trains of 70 and 100 dB-peSPL amplitudes. ABR W-I and W-V amplitudes were defined as half the peak-to-trough amplitude and calculated between a positive peak and subsequent negative trough (see Picton, 2010).

### Simulated and recorded EFRs: time domain comparison

To investigate whether the EFR, as a population response across neurons of different CFs, follows the on-CF ANF response trends, Fig. 1i-l shows simulated (open traces) and recorded grand-average EFR waveforms per group (solid traces). Simulated and recorded NH EFRs (Fig. 1i-l, green) generally followed the trends observed in the ANF responses by showing the strongest response maxima for the RAM25 stimulus. Comparing the H10AM and RAM25 EFRs shows that long IPIs and short duty cycles are important, but not sufficient, to evoke a strong EFR as low-level stimulation between the envelope maxima (H10AM) can end up reducing the EFR. It is noteworthy that the RAM50 recording showed two peaks per cycle whereas neither the RAM50 simulations, nor the RAM25 responses showed such a double response peak within an envelope cycle. Our simulations showed that double responses can occur when apical off-CF BM vibrations (approximately up to one octave below the 4-kHz CF) contribute to the ANF population response. However, the CN/IC filtering properties of our model removed this second peak from the EFR simulations.

In line with the ANF simulations, synaptopathy reduced the simulated peak-to-peak EFR amplitudes. When comparing the simulated EFR declines to our recordings, we see that EFRs of both older listener groups (oNH and oHI) were reduced compared to the yNH group. Our simulations suggest that the range of experimentally-observed EFR reductions could be explained by the synaptopathy aspect of SNHL, as several of the stimulation paradigms were shown not to be affected by much cochlear gain loss. Following this line of thought, we predict that the oHI listeners had a stronger degree of synaptopathy because their RAM EFRs were smaller than those of the oNH group. Lastly, it is noteworthy that the response difference between oNH and oHI groups disappeared for the H10AM condition (Fig. 1l). This observation corroborates the simulations showing that the OHC-damage aspect can counteract the synaptopathy-induced response reduction for the H10AM, but not RAM, stimuli.

### Group EFR magnitudes across stimulus conditions

Figure 5a compares recorded (filled symbols) and simulated (triangles) EFR magnitudes across conditions and subject groups, and Table A2 summarizes the group means and interquartile ranges. Recorded NH EFR magnitudes were significantly smaller for the SAM stimulus than for the other stimuli: paired t-tests between SAM and the other conditions showed p*<*0.001 for all tests. In further agreement with the simulations, the longer duty cycle RAM50 stimuli yielded significantly smaller EFR magnitudes than the RAM25 stimulus (NH ptp/rms condition, t=6.87/8.65, p*<*0.001). Despite the sensitivity of the SAM-EFR metric to synaptopathy (model simulations and Parthasarathy et al., 2018), the data points showed considerable overlap across the three subject groups and this limits the potential of this marker for diagnostic purposes. In contrast, NH EFR magnitudes to the other stimuli were overall larger and showed a greater spread around the mean. Individual differences were emphasized and this could benefit diagnostic interpretation. Specifically, the RAM25 magnitudes showed non-overlapping interquartile ranges between the groups (see also Table A2). Within the cohort of yNH and oNH listeners, the RAM25-EFR showed a considerable oNH group decline (t=4.91, p*<*0.001), whereas the SAM-EFR did not (t=1.3, p=0.1). Because experimental RAM25-EFRs were overall 3 times stronger than SAM EFRs, individual responses to the latter metric would more quickly reach the experimental noise floor or be immersed into the standard deviations of the metrics, which were statistically similar for both response types (t=0.89, p=0.37, N=44, EFR standard deviations of SAM vs RAMptp). The observed age-related EFR decline in the RAM (but not SAM) condition, along with how the EFR magnitudes changed across conditions is entirely consistent with how the same degree of simulated synaptopathy yielded an equal % reduction in EFR strength (upto 81% for the most severe simulated synaptopathy pattern).

We argue that the age-related EFR decline observed for oNH listeners, followed by a further EFR reduction for oHI listeners to the RAM25-EFR, is not due to audiometric threshold or age-differences between the groups, but rather reflects increasing degrees of synaptopathy, as predicted by our model simulations. Even though we cannot directly prove this without histopathology, there are a number of observations that support our conclusion: age-differences between the oNH and HI group could not be responsible for their group differences as the groups were age-matched. In contrast, threshold differences between the groups might explain the lower EFR magnitudes for the oHI than oNH group, but this is unlikely for two reasons: (i) our EFR simulations in Fig. 5 show that synaptopathy has a much greater effect on the EFR (upto 81% reduction) than possibly co-existing OHC damage (5-10% change depending on the stimulus). (ii) A second, experimental argument in favor of a synaptopathy explanation relates to the observation that RAM-EFR group-mean differences were larger between the yNH and oNH/oHI groups (t=4.91 and 8.38 with p*<*0.001 in both cases) than between the oNH and oHI group (t=3.70 and p=0.001), whereas the 4-kHz hearing thresh-old differences between the groups (Fig. 2) were larger between the oNH and oHI group (t=12.48, p*<*0.001) than between the yNH and oNH group (t=6.04; *<*0.001). Comparing the recordings and model simulations directly, and in the presumed absence of other, non-SNHL related, age-effects on the EFR, the oNH group had EFRs corresponding to a simulated 0L-0M-7H SR synpatopathy profile, whereas the oHI group most closely matched the 0L-0M-3H SR profile.

### Individual EFR differences: roles of hearing threshold and age

To further explore the respective roles of hearing sensitivity and age, Fig. 6 depicts the relationship between individual EFR magnitudes and 4-kHz DPOAE thresholds, and shows mean EFR magnitudes for groups separated by their 4-kHz audiometric threshold (yNH/oNH *<* 25 dB, Table A1) or age (yNH *<* 30 y/o and oNH/oHI *>* 60 y/o). The group data on top of Fig. 6a shows that it was not possible to discriminate between younger and older participants on the basis of the SAM-EFR magnitude. The same conclusion is drawn for the normal and elevated audiometric threshold groups. The weaker SAM-EFR responses thus appeared affected by both an age and OHC-damage aspect of SNHL. Despite overall stronger H10AM EFR magnitudes (Fig. 6b), they were similarly unable to segregate between listener groups of young/old age, or groups with normal/elevated thresholds. In contrast, the RAM-EFR magnitudes (Fig. 6c,d) were able to separate listeners into groups of younger and older listeners, demonstrating that this condition was more susceptible to the age-related aspect of SNHL than to OHC damage. At the same time, the separation into different age groups was better on the basis of the RAM-EFR than on the basis of the DPOAE threshold. Even though OHC and age-related SNHL deficits likely coexist in older listeners, the RAM stimulus was better able to isolate the age-related aspect than the other considered stimuli. These results corroborate our model simulations which illustrate how on-CF ANF responses show a near-differential sensitivity to synaptopathy for the RAM stimuli, and a mixed sensitivity to OHC damage and synaptopathy for the SAM and H10AM stimuli (Fig. 1). Simulated EFRs (Fig. 5) followed this trend by showing a stronger sensitivity of the RAM EFR to synaptopathy, i.e., an absolute EFR reduction upto 0.12 *µ*V (RAM25) vs 0.08 *µ*V for SAM, while possible confounding effects of OHC damage were 10 times smaller for the RAM25 condition (0.014 *µ*V). Because animal studies of age-related, and histologically-verified, synaptopathy show reduced EFRs (Sergeyenko et al., 2013; Fernandez et al., 2015; Möhrle et al., 2016b; Parthasarathy & Kujawa, 2018), we have strong circumstantial evidence that the RAM-EFR magnitude was successful at separating listeners into groups with and without age-related auditory de-afferentation, including cochlear synaptopathy.

**Figure 6.**
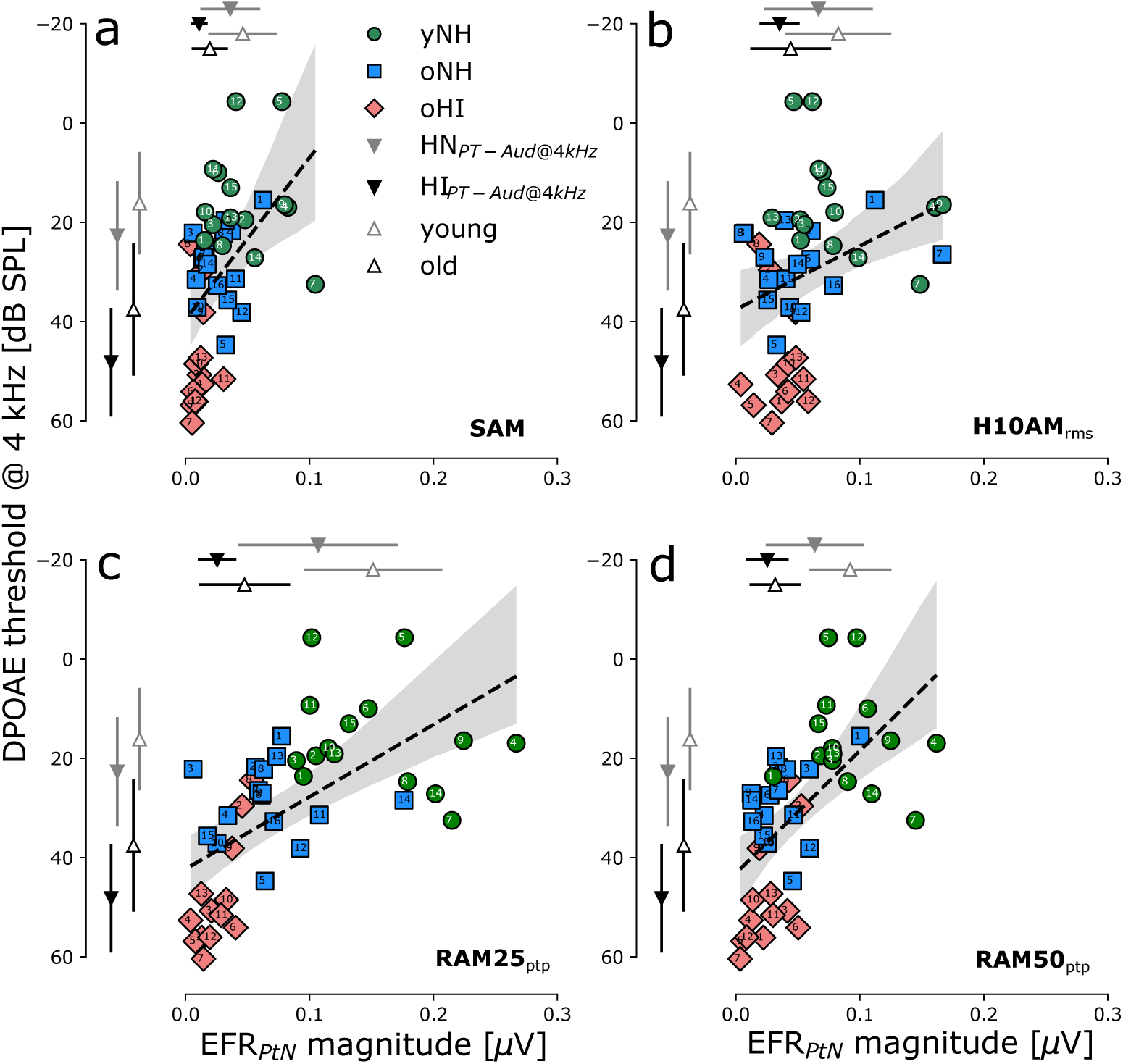
Linear regression plots for EFR magnitudes evoked by AM stimuli with different envelope shapes and 4-kHz DPOAE thresholds. Top and left error bars indicate groups means and standard deviations of normal (yNH&oNH) and elevated (oHI) audiometric threshold (at 4 kHz) groups (downward triangles), as well as young (yNH) and older (oNH&oHI) groups (upward triangles).

To further explore the relationship between age-related SNHL factors (e.g. synaptopathy and OHC damage) and individual EFR magnitudes, we constructed a linear regression model of the form: EFR_PtN_ = *β*_0_ + *β*_1_ *·* Age + *β*_2_ *·* DPOAE_@4kHz_ which included data of all participants (N=44), and decomposed the models’ R^2^ into commonality coefficients (using R Core Team, 2019; Nimon et al., 2008). Table 1 shows that approximately half the total explained variance (common) was attributed to significant relationships between the EFR magnitude and predictor variables age and DPOAE_@4kHz_ across conditions. However, OHC damage (as reflected in the DPOAE_@4kHz_ threshold) showed the smallest contribution to the EFR magnitude for all conditions. When the variance of age-related factors were accounted for, the unique DPOAE_@4kHz_ contribution became negligible. This commonality analysis shows that –in case of co-occurring age-related and OHC-damage SNHL aspects–, the observed relationships between the EFR magnitude and DPOAE_@4kHz_ were merely driven by the age-related aspect. Supported by Table 1: Results for the multiple regression model and commonality analysis the model simulations, we believe that age-induced synaptopathy was responsible for driving these regression model outcomes.

**Table 1:** Results for the multiple regression model and commonality analysis.

| Stimulus | R<sup>2</sup> | Adj. R<sup>2</sup> | p-value | beta | unique<sup>†</sup> | common<sup>†</sup> |
| --- | --- | --- | --- | --- | --- | --- |
| SAM | 0.314 | 0.281 | 0.0004 | $\beta_1=-0.0005^*$ | 26.60 | 64.47 |
| | | | | $\beta_2=-0.0003$ | 8.93 | |
| H10AM<sub>rms</sub> | 0.220 | 0.182 | 0.0061 | $\beta_1=-0.0010^*$ | 52.89 | 47.01 |
| | | | | $\beta_2=-0.0001$ | 0.10 | |
| RAM50<sub>ptp</sub> | 0.606 | 0.587 | <0.0001 | $\beta_1=-0.0013^{***}$ | 41.98 | 56.06 |
| | | | | $\beta_2=-0.0004$ | 1.96 | |
| RAM25<sub>ptp</sub> | 0.585 | 0.565 | <0.0001 | $\beta_1=-0.0022^{***}$ | 38.18 | 58.64 |
| | | | | $\beta_2=-0.0008$ | 3.18 | |

Significance codes: $p < 0.0001$ \*\*\*, $p < 0.001$ \*\*, $p < 0.01$ \*, $p < 0.05$ .
<sup>†</sup> % of R<sup>2</sup>

| Subj# | S | A | 0.125 | 0.25 | 0.5 | 0.75 | 1 | 1.5 | 2 | 3 | 4 | 6 | 8 | PTA |
| --- | --- | --- | --- | --- | --- | --- | --- | --- | --- | --- | --- | --- | --- | --- |
| Young participants with normal hearing (yNH) |  |  |  |  |  |  |  |  |  |  |  |  |  |  |
| 1 | m | 27 | 5 | 5 | 10 | 5 | -5 | 5 | 0 | 5 | **0** | 10 | 5 | 4.1 |
| 2 | f | 27 | 5 | 10 | 10 | 10 | 10 | 0 | 5 | 0 | **0** | 0 | 0 | 4.5 |
| 3 | m | 25 | 0 | 0 | 0 | 0 | 0 | 0 | 0 | 0 | **5** | 0 | 5 | 0.9 |
| 4 | f | 22 | 0 | 0 | 5 | 5 | 0 | 0 | 5 | 5 | **5** | 5 | 10 | 3.6 |
| 5 | f | 27 | 5 | 0 | 0 | 0 | 0 | 0 | 0 | 0 | **0** | 5 | 5 | 1.4 |
| 6 | f | 20 | 0 | -5 | 0 | 0 | 0 | 5 | 0 | 5 | **0** | 0 | 0 | 0.5 |
| 7 | f | 24 | 5 | 5 | 5 | 0 | 5 | 0 | 0 | 0 | **0** | 0 | 0 | 1.8 |
| 8 | f | 25 | 0 | 0 | 0 | 0 | 0 | 0 | 0 | 5 | **10** | 5 | 10 | 2.7 |
| 9 | f | 28 | 0 | 0 | 0 | 0 | 0 | 5 | 5 | 0 | **5** | 0 | 0 | 1.4 |
| 10 | f | 23 | 0 | 0 | 5 | 0 | 0 | 0 | 0 | 0 | **5** | 5 | 10 | 2.3 |
| 11 | m | 22 | 0 | 0 | 0 | 0 | 5 | 5 | 0 | 0 | **0** | 0 | 0 | 0.9 |
| 12 | m | 24 | 5 | 5 | 0 | 5 | 0 | 0 | 0 | 0 | **5** | 5 | 0 | 2.3 |
| 13 | m | 25 | 5 | 5 | 5 | 10 | 10 | 5 | 5 | 15 | **5** | 5 | 15 | 7.7 |
| 14 | m | 23 | 10 | 15 | 15 | 15 | 15 | 10 | 10 | 10 | **10** | 10 | 15 | 12.3 |
| 15 | m | 26 | 5 | 5 | 10 | 10 | 5 | 5 | 0 | 5 | **0** | 0 | 5 | 4.5 |
| Older participants with normal hearing(oNH) |  |  |  |  |  |  |  |  |  |  |  |  |  |  |
| 1 | f | 63 | 15 | 10 | 5 | 10 | 5 | 15 | 5 | 0 | **0** | 25 | 30 | 10.9 |
| 2 | f | 61 | 10 | 10 | 10 | 5 | 5 | 5 | 5 | 10 | **10** | 15 | 45 | 11.8 |
| 3 | m | 63 | 5 | 5 | 10 | 15 | 15 | 5 | 10 | 10 | **15** | 45 | 65 | 18.2 |
| 4 | m | 62 | 5 | 5 | 5 | 10 | 5 | 5 | 5 | 5 | **10** | 10 | 15 | 7.3 |
| 5 | f | 65 | 10 | 10 | 15 | 15 | 15 | 10 | 10 | 20 | **10** | 15 | 30 | 14.5 |
| 6 | m | 64 | 0 | 0 | 0 | 0 | 5 | 5 | 5 | 5 | **10** | 10 | 35 | 6.8 |
| 7 | f | 62 | 10 | 10 | 15 | 15 | 20 | 10 | 10 | 10 | **15** | 30 | 30 | 15.9 |
| 8 | m | 65 | 5 | 5 | 5 | 5 | 5 | 5 | 0 | 10 | **10** | 30 | 35 | 10.5 |
| 9 | m | 65 | 10 | 5 | 5 | 5 | 5 | 10 | 5 | 15 | **10** | 25 | 50 | 13.2 |
| 10 | f | 66 | 5 | 10 | 10 | 10 | 15 | 20 | 15 | 20 | **15** | 30 | 55 | 18.6 |
| 11 | m | 66 | 20 | 20 | 20 | 20 | 15 | 15 | 15 | 15 | **15** | 40 | 45 | 21.8 |
| 12 | f | 65 | 20 | 20 | 20 | 25 | 25 | 20 | 20 | 20 | **15** | 30 | 30 | 22.3 |
| 13 | f | 62 | 0 | 0 | 0 | 10 | 10 | 10 | 5 | 5 | **10** | 10 | 15 | 6.8 |
| 14 | m | 67 | 15 | 5 | 5 | 10 | 10 | 5 | 20 | 15 | **15** | 15 | 45 | 14.5 |
| 15 | m | 67 | 10 | 5 | 5 | 5 | 5 | 5 | 5 | 10 | **10** | 15 | 25 | 9.1 |
| 16 | f | 65 | 10 | 10 | 5 | 5 | 20 | 10 | 10 | 10 | **15** | 20 | 25 | 12.7 |
| Older participants with elevated hearing thresholds (oHI) |  |  |  |  |  |  |  |  |  |  |  |  |  |  |
| 1 | f | 64 | 0 | 5 | 10 | 10 | 10 | 10 | 20 | 25 | **40** | 35 | 35 | 18.2 |
| 2 | f | 67 | 20 | 20 | 25 | 30 | 35 | 30 | 30 | 35 | **25** | 50 | 60 | 32.7 |
| 3 | f | 64 | 0 | 5 | 10 | 15 | 10 | 25 | 25 | 35 | **40** | 50 | 65 | 25.5 |
| 4 | m | 66 | 5 | 5 | 10 | 10 | 15 | 15 | 15 | 25 | **35** | 40 | 60 | 21.4 |
| 5 | m | 64 | 5 | 5 | 10 | 15 | 20 | 25 | 25 | 25 | **40** | 45 | 50 | 24.1 |
| 6 | f | 66 | 10 | 10 | 10 | 10 | 5 | 5 | 5 | 25 | **35** | 40 | 40 | 17.7 |
| 7 | f | 66 | 10 | 10 | 10 | 25 | 20 | 25 | 35 | 35 | **50** | 55 | 60 | 30.5 |
| 8 | f | 61 | 10 | 15 | 20 | 20 | 15 | 25 | 30 | 30 | **30** | 35 | 35 | 24.1 |
| 9 | m | 65 | 20 | 20 | 20 | 20 | 20 | 20 | 25 | 35 | **45** | 60 | 65 | 31.8 |
| 11 | m | 64 | 5 | 10 | 10 | 20 | 25 | 15 | 10 | 25 | **40** | 60 | 60 | 25.5 |
| 12 | m | 66 | 5 | 0 | 5 | 5 | 5 | 5 | 5 | 25 | **30** | 60 | 70 | 19.5 |
| 13 | f | 68 | 15 | 15 | 20 | 20 | 20 | 30 | 25 | 35 | **40** | 35 | 60 | 28.6 |
| 14 | f | 67 | 5 | 5 | 5 | 5 | 5 | 10 | 10 | 10 | **40** | 55 | 55 | 18.6 |

**Table 3:**
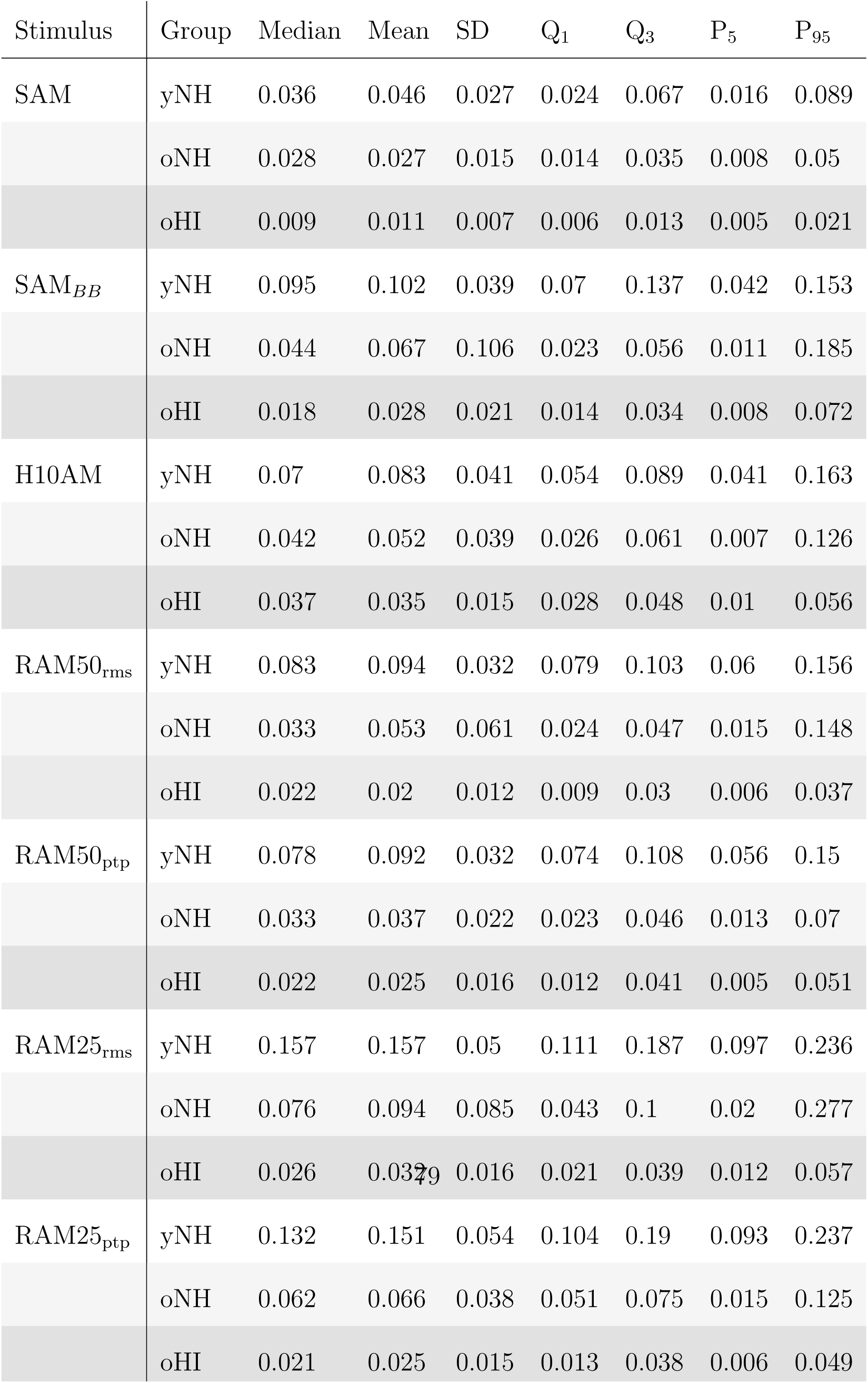
**Table A2:** Median, mean, standard deviation (SD), 1^st^ and 3^rd^ quartile (Q), 5^th^ and 95^th^ percentile (P) of the EFR magnitudes (in *µ*V) evoked by different stimuli.

**Table 4:** **Table A3:** Median, mean, standard deviation (SD), 1^st^ and 3^rd^ quartile (Q), 5^th^ and 95^th^ percentile (P) of the ABR amplitudes (in *µ*V) evoked by different stimuli.

| Stimulus | Group | Median | Mean | SD | Q<sub>1</sub> | Q<sub>3</sub> | P<sub>5</sub> | P<sub>95</sub> |
| --- | --- | --- | --- | --- | --- | --- | --- | --- |
| ABR W-I<sup>70†</sup> | yNH | 0.079 | 0.081 | 0.034 | 0.061 | 0.09 | 0.04 | 0.131 |
|  | oNH | 0.044 | 0.053 | 0.025 | 0.032 | 0.074 | 0.023 | 0.086 |
|  | oHI | 0.018 | 0.018 | 0.011 | 0.013 | 0.023 | 0.009 | 0.028 |
| ABR W-I<sup>100†</sup> | yNH | 0.126 | 0.134 | 0.043 | 0.112 | 0.165 | 0.063 | 0.194 |
|  | oNH | 0.093 | 0.083 | 0.046 | 0.043 | 0.113 | 0.022 | 0.15 |
|  | oHI | 0.072 | 0.07 | 0.032 | 0.043 | 0.08 | 0.028 | 0.12 |
| ABR W-V<sup>70†</sup> | yNH | 0.153 | 0.176 | 0.07 | 0.131 | 0.211 | 0.089 | 0.276 |
|  | oNH | 0.077 | 0.093 | 0.055 | 0.062 | 0.125 | 0.025 | 0.178 |
|  | oHI | 0.065 | 0.071 | 0.043 | 0.04 | 0.099 | 0.019 | 0.135 |
| ABR W-V<sup>100†</sup> | yNH | 0.152 | 0.179 | 0.083 | 0.141 | 0.181 | 0.111 | 0.331 |
|  | oNH | 0.123 | 0.149 | 0.091 | 0.098 | 0.179 | 0.06 | 0.294 |
|  | oHI | 0.115 | 0.111 | 0.039 | 0.08 | 0.12 | 0.062 | 0.169 |
| ABR W-V<sup>100†</sup><sub>(P5-P3)</sub> | yNH | 0.267 | 0.271 | 0.067 | 0.233 | 0.314 | 0.167 | 0.371 |
|  | oNH | 0.213 | 0.201 | 0.062 | 0.153 | 0.233 | 0.119 | 0.313 |
|  | oHI | 0.165 | 0.169 | 0.061 | 0.118 | 0.207 | 0.091 | 0.265 |

† dB peSPL

Lastly, we investigated whether the adopted EFR analysis method had an effect on the regression model outcomes. To this end, we also constructed a multiple regression model with the commonly-adopted SAM-EFR magnitude metric, i.e. the spectral peak F_n@120 Hz_ (Fig. 4), as the explaining variable. In contrast to using our EFR markers (Eq. 5), this model did not reach significance (multiple R^2^ = 0.073, adjusted R^2^ = 0.028, p-value = 0.2127). The weaker SAM-EFR response, inter-subject variability in the noise floor (NF), and overall underestimation of stimulus-envelope related energy in the EEG spectrum (i.e. due to the omission of harmonics), were together responsible for this outcome and stress the need for optimized stimuli and analysis methods.

### Comparison between EFR magnitudes and ABR amplitudes

Figure 5b compares SAM and RAM25 EFRs to features derived from low-rate (10 Hz) ABRs recorded in the same listeners. ABR amplitudes were defined as half the peak-to-trough amplitude, to allow a fair comparison to the EFR magnitude as calculated using Eq. 5. NH ABR amplitudes followed the expected trends observed in normative data (Picton 2010), with overall smaller W-I than W-V amplitudes and larger amplitudes for higher SPL (see also Table A3). Both oNH and oHI groups had reduced group median W-I amplitudes compared to the yNH group. However, the interquartile ranges overlapped and only showed significant differences between yNH and oHI W-I amplitudes for the 70-dB-peSPL condition (t = 4.48, p = 0.02). The 100-dB-peSPL W-I amplitude was able to separate the yNH and oNH/oHI groups (t=3.07, p=0.004 and t=4.31, p=0), but not the oNH and oHI groups, consistent with the view that this marker was able to detect age-related SNHL aspects on a group level. ABR W-V amplitudes were overall larger, but nevertheless showed considerable overlap between the groups, pointing to a limited diagnostic sensitivity in detecting age or OHC-damage aspects of SNHL. The 70-dB-peSPL W-V amplitude separated the yNH from the oNH/oHI groups (t=3.48, p= 0.002 and t=4.6, p*<*0.001), and the 100-dB-peSPL W-V condition separated the yNH from the oHI group (t=2.76, p=0.01), but not the yNH from the oNH group.

Comparing ABRs to EFRs recorded from the same listeners revealed that EFRs had reduced magnitude distributions compared to the ABRs (e.g. compare the interquartile ranges for the oNH and oHI groups, resp.). EFRs were computed using automatic procedures, and were corrected for by the individual NF, which may partly explain this observation. Despite similar RAM25-EFR and ABR amplitudes, the EFR group means were much more separated than the ABR group means. This can be explained by their differential sensitivity to different aspects of SNHL. A recent modeling study showed that both synaptopathy and OHC-damage reduce the generator strength of the ABR (Verhulst et al., 2016). The present study shows that simulated RAM-EFRs can be altered by 4-10 % due to OHC loss, while synaptopathy can reduce its strength by 81%. Taken together, we conclude that the RAM25-EFR has a better sensitivity than the ABR in isolating the age-related (synaptopathy) aspect of SNHL.

### Effect of IPI duration and stimulation rate on the EFR

To better understand how the stimulus envelope shape affects the EFR sensitivity to different SNHL aspects, and to optimize the stimulation paradigms for diagnostic purposes, we conducted an additional number of simulations in which we modified the duty cycle and IPI (Fig. 7) or modulation rate (Fig. 8) of the RAM stimuli. Because stimulation with short-duty-cycle RAM stimuli resembles ABR stimulation paradigms, we investigated whether there was a benefit of adopting an EFR paradigm (response to modulation rate) over an ABR paradigm (onset response peak). To this end, we simulated how the interplay between RAM plateau duration and IPI duration affected the EFR magnitude. In Fig. 7a, we considered stimulation with a modulation rate of 10 Hz (akin the click-ABR rate), while changing the duty cycle from 0.2 to 25%. The 25% condition corresponded to the longest plateau duration and the 0.2 % condition resembled classical ABR stimulation paradigms most closely. We changed the IPI of the reference 120-Hz RAM EFR condition to 100 ms to simulate the 10-Hz modulation rates (i.e., the plateau durations were the same in both 10 and 120 Hz conditions while the IPI was different).

**Figure 7.**
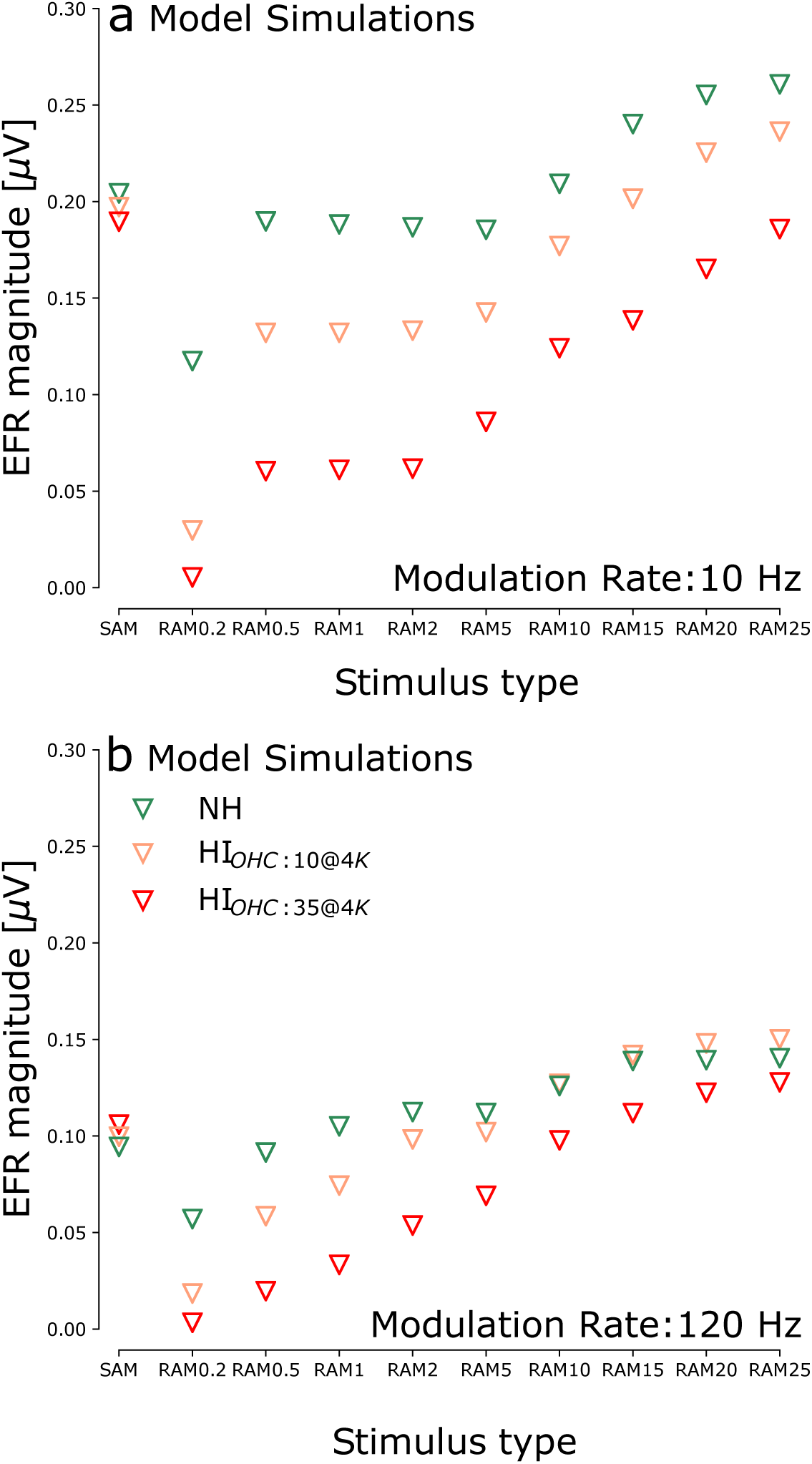
Simulated EFR magnitudes for 10 Hz (**a**) and 120 Hz (**b**) modulation rate RAM stimuli with the same peak-to-peak amplitude as the reference 70-dB-SPL SAM tone. EFR magnitudes are shown for the NH model as well as for models with sloping cochlear gain loss (HI_OHC:10@4K_ and HI_OHC:35@4K_).

**Figure 8.**
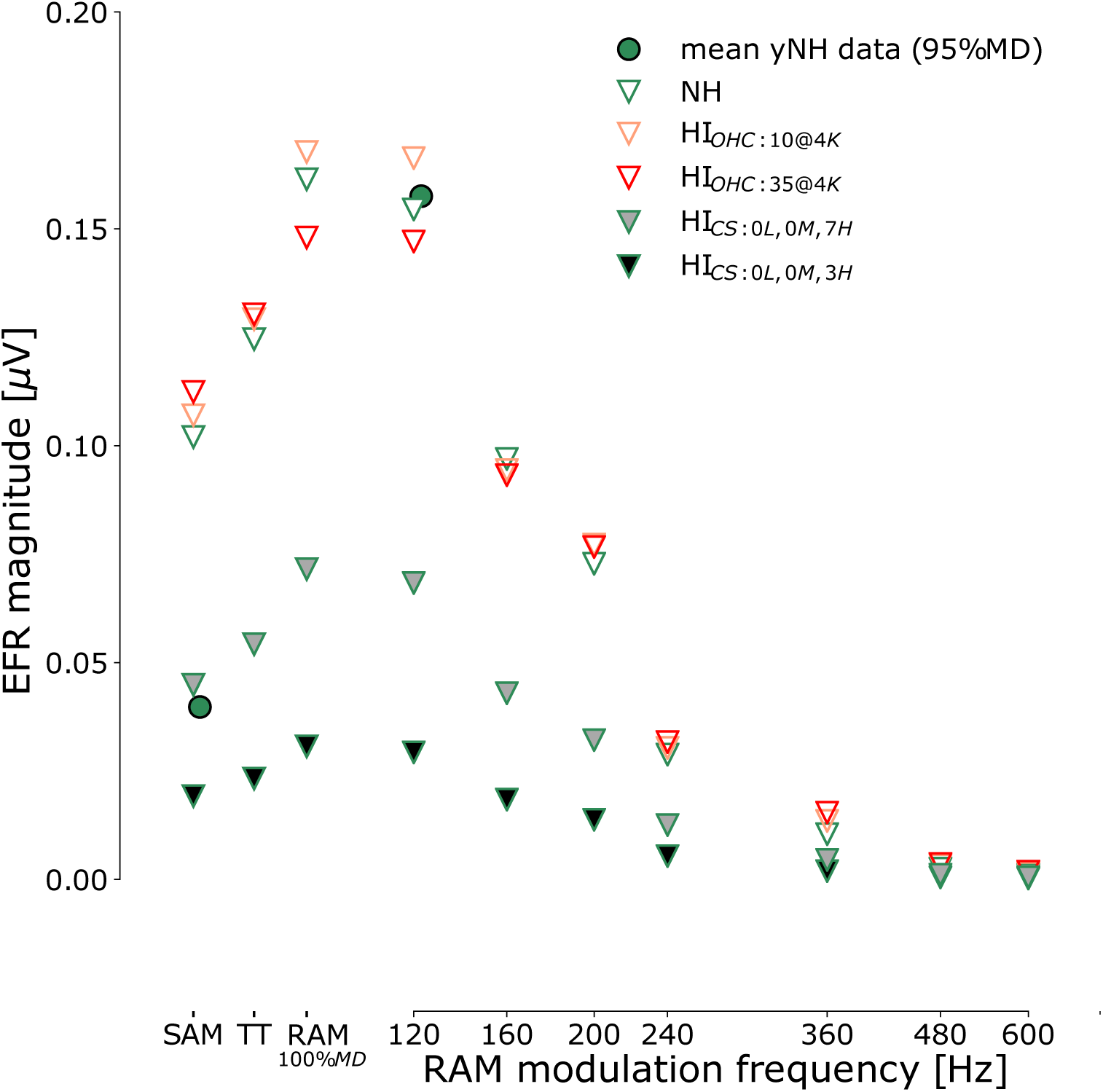
Simulated EFR magnitudes for reference SAM, transposed tone (TT; van de Par & Kohlrausch, 1997) and RAM stimuli with a duty cycle of 25%. Stimulus levels were 70 dB SPL for all conditions and mean experimental EFR magnitudes for the yNH group are also shown. The modulation frequency was 120 Hz for the SAM and TT conditions, and was varied for the RAM EFR simulations. All modulation depths were 95%, except for the RAM_100%_*_MD_* condition. Simulations are shown for the NH model (green triangles), as well as for models with simulated cochlear gain loss (salmon, red triangles) and cochlear synaptopathy (gray filled triangles).

In agreement with how ABRs to low-repetition (10 Hz) click trains yield robust ABRs (Picton, 2010), simulated 10-Hz RAM EFRs evoked overall stronger responses than the 120-Hz condition (Fig. 7a vs b). At the same time, stimuli of both repetition rates evoked substantially reduced EFRs when the duty cycle reduced from 25% to 0.2%. These simulations suggest that one of the explaining factors for weak EFRs could relate to the lower amount of sound energy carried in each short duty cycle, resulting in a compromised synchronous ANF response. Moreover, large duty cycles (e.g. 50% as used in RAM50) were also shown to evoke reduced EFRs when compared to the 25% duty cycle (Fig. 1g,k vs f,j). The silence interval between the stimulus peaks (IPI) appeared hence more important than the duration of the duty cycle to yield robust EFRs. However, too short, or too long, plateau durations can also compromize the stimulation efficiency, and hence point to a sweet-spot duty cycle of 15-25% for most-efficient EFR stimulation.

OHC-damage affected the simulated EFRs differently depending on the duty cycle: stimuli with short (click-like) duty cycles evoked responses that were strongly influenced by the OHC aspect of SNHL, and this effect reduced as the duty-cycle increased. Secondly, the overall stimulation rate also matters: the 10-Hz condition was more affected by cochlear gain loss than the 120-Hz condition. The reason for these differences was, in the model, attributed to ANF responses which were strongly affected by the active components of the BM impulse response when the stimulation rate was low. Consequently, a strongly attenuated response was observed when OHC dysfunction was simulated. In contrast, the 120-Hz AM stimuli evoked more-saturated ANF responses, yielding overall lower NH responses which were less impacted by OHC dysfunction. To support the predominantly saturated ANF response origin of the 120-Hz condition, simulated EFRs (both SAM and RAM25) with duty cycles greater than 10% showed slightly elevated EFR magnitudes for high-frequency OHC damage (compare NH to HI_OHC:10@4K_ in Fig. 7b). In the model, these enhanced EFR magnitudes stemmed from the linearized (basal) cochlear responses and attenuated BM input to the ANF in the IPI, which pulls the ANFs out of saturation to yield a stronger modulated response (Joris & Yin, 1992). From the OHC-damage simulations in Fig. 7, and prior simulations in Fig. 1e-l and Fig. 5a regarding the sensitivity of the 120-Hz RAM EFRs to the number of intact ANFs, we conclude that faster-rate EFRs, with duty cycles between 15-25%, are most optimal to quantify the synaptopathy aspect of SNHL, even when OHC-damage is concomitantly present.

### EFRs to transposed-tone, SAM or RAM stimuli

Whereas animal studies of SNHL and supra-threshold hearing often use SAM tones (Zhong et al., 2014; Shaheen et al., 2015; Möhrle et al., 2016b; Parthasarathy et al., 2018), human studies have either considered SAM stimuli (e.g. Goossens et al., 2016; Paul et al., 2017a,b; Garrett & Verhulst, 2019), transposed tones (e.g. Bharadwaj et al., 2015; Prendergast et al., 2017a; Guest et al., 2017), or responses to different AM envelope shapes (e.g. John et al., 2002; Van Canneyt et al., 2019). As our simulations show, these stimulus details may be important to consider, as envelope modifications can both influence the response strength and their sensitivity to different SNHL aspects. In agreement with the Van Canneyt et al. (2019) study, our simulations show that short duty cycles and steeply rising envelope shapes yield more synchronized ANF responses and stronger EFRs. Our observations also corroborate the experimental findings of Dreyer & Delgutte (2006) showing stronger ANF responses for stimuli with more sharply rising envelopes (Fig. 1). To study the expected effects for a set of commonly used stimuli, Fig. 8 compares EFR simulations to 120-Hz modulated SAM, RAM25, and transposed tone (TT) stimuli (van de Par & Kohlrausch, 1997). The NH EFR magnitudes increased from SAM to TT to RAM25 stimulation, following their gradual steeper envelope shapes and increasing IPIs. In this comparison, it is important to bear in mind that the experimental difference between the SAM and RAM25 response was much greater than for the model simulations. Even though Fig. 5 showed that simulated yNH EFRs followed the absolute magnitudes and predicted changes to stimulus conditions well, the SAM_PT_ EFR was an outlier. We noticed a contribution of very basal, off-CF AM contributions to the SAM_PT_ EFR in the model, which was not observed for the other stimulus conditions. In future simulations, we could mask these very basal off-CF contributions with a neural noise-floor, but we refrained from doing this here. Such very basal off-CF AM contributions did not affect the other conditions, even the simulated SAM_BB_ response corresponded nicely to the experimental findings. However, as a cautionary note for the SAM_PT_ EFR: the recordings should bear more weight towards our conclusions than the simulations.

Going deeper into the EFR generator mechanisms, we studied whether stronger RAM vs SAM EFR magnitudes were caused by a broader tonotopic excitation of the RAM stimulus, or whether the shape of the temporal envelope was responsible for eliciting a stronger and more synchronized neuronal response. To this end, we compared RAM and SAM EFRs to EFRs evoked by a broadband white noise carrier (Fig. 5a; SAM_BB_). SAM_BB_ magnitudes were generally larger than SAM_PT_ magnitudes (NH: t=6.61, p*<*0.001), but smaller than RAM25 EFRs (NH: t=7.25, p*<*0.001; see also Table A2). We hence conclude that the shape of the temporal envelope was more important than the bandwidth of the carrier to evoke a strong EFR.

Studying the effect of SNHL, Fig. 8 shows that all three commonly used EFR types were highly sensitive to synaptopathy (i.e. gray filled symbols), whereas OHC damage impacted them differently: both SAM and TT EFRs showed slightly stronger EFRs, whereas the RAM EFR showed a small reduction for the highest degree of SNHL. The SAM EFR enhancements resulted from stronger on-CF single-unit peak firing rates, whereas these remained unaffected for the RAM EFR (see Fig. 1e-f). The OHC-damage-related EFR reduction in the RAM condition was instead caused by a reduction of off-CF AM contributions in the tails of the cochlear excitation patterns, a mechanism earlier discussed in Encina-Llamas et al. (2019); Keshishzadeh et al. (2019, 2020). When using a single modulation depth and modulation fre-quency, we can conclude that none of the considered EFR stimulus paradigm was 100% insensitive to OHC damage. However, the RAM stimulus had a clear advantage in that it did not enhance the on-CF ANF response, whereas the SAM and TT stimuli did. Aside from the on-CF contributions, reduced off-CF AM contributions to the EFR caused by OHC damage are greater as the stimulus envelope is modified from SAM to TT to RAM stimulation which cause increasingly wider cochlear excitation patters (even for tonal carriers). The relative contribution of these two –on and off-CF– EFR sources caused a net increase for the SAM and TT condition, whereas it caused a net decrease for the RAM condition when there was considerable OHC damage. In summary, the % EFR reduction (i.e., normalized to the NH EFR) was −10, −4 and +4 % for the 35 dB HL OHC-damage pattern, whereas it was 81 % for the most severe, 0L,0M,3H synaptopathy pattern. In absolute EFR values, the RAM EFR was reduced by 0.08, 0.10 and 0.12 *µ*V resp. for the same degree of synaptopathy, demonstrating the superior sensitivity of the RAM EFR metric for diagnostic purposes.

Lastly, to further optimize the RAM stimulation paradigm, we investigated whether further increasing the modulation depth from 95% to 100% would benefit the EFR and its sensitivity to SNHL. The added 100% MD condition in Fig. 8 shows that a further EFR magnitude increase is expected, while similar SNH-effects are predicted. The 100% MD condition is hence recommended when conduction new experiments with the RAM stimuli.

### Effect of envelope modulation frequency on the EFR

EFRs evoked by AM signals are known to vary in magnitude depending on the modulation rate (Purcell et al., 2004; Parthasarathy et al., 2018). Our recording paradigm adopted a rate of 120 Hz to focus on subcortical EFR generators (Purcell et al., 2004; Herdman et al., 2002; Bidelman, 2015, 2018), and in Fig. 8 we studied whether further increasing the modulation frequency might emphasize more peripheral sources. However, we should consider the strength of the recorded signal as well when increasing the modulation rate. Previous studies have observed EFR magnitude declines for modulation rates above 60-70 Hz, that often become statistically indistinguishable from the background noise for modulation rates above 250 Hz (Purcell et al., 2004; Picton, 2010; Garrett & Verhulst, 2019). Figure 8 depicts simulated RAM25 EFR magnitudes for modulation frequencies between 120 and 600 Hz using a fixed 95% modulation depth. EFR magnitudes became less sensitive to OHC deficits as the modulation frequency increased to higher rates, e.g. at 240 Hz there was no influence of OHC damage. However, this increased differential sensitivity to synaptopathy occurred at the cost of overall reduced EFR magnitudes, which might compromise the sensitivity of the metric toward detecting individual differences.

## Discussion

We adopted a combined computational and experimental approach to investigate which EFR paradigms and analysis methods would enhance its sensitivity to isolate the synaptopathy aspect of SNHL in listeners who may have mixed OHC-damage/synaptopathy pathologies. The modeling work incorporated our latest knowledge of the physiology of hearing and hearing damage, and predicted the outcomes of the experimental study well. Even though we did not have access to animal physiology methods in this study, our approach strongly supports the use of the RAM25 stimulus as a sensitive marker for synaptopathy in humans.

### Stimulus envelope encoding in the aging auditory system

Aside from our main pursuit to develop more sensitive EFR paradigms for synaptopathy diagnosis, the outcomes of this study shed light on how SNHL affects envelope encoding in the aging auditory system. Particularly, Fig. 5 shows that envelope coding in the older groups is significantly different when SAM or RAM stimuli are used, and this result is entirely attributed to the overall stronger EFR responses in the RAM conditions which emphasized individual response differences. If we were to base our conclusions solely on the SAM results, we would conclude that aging (with clinically normal thresholds upto 4 kHz) does not significantly impact envelope coding, whereas the conclusion is drastically different for the RAM condition that does show a significant decline. The improved EFR sensitivity in the RAM EFR not only exacerbated the group differences between the yNH, oNH and oHI groups, our model simulations suggest that these differences could be explained by synaptopathy. The RAM25 condition, for which simulated and recorded EFR magnitudes corresponded well, predicts a mean 85% ANF deafferentation for the oHI listeners, and a 63% ANF de-afferentation for the oNH listeners. Our predicted degree of de-afferentation for the oNH group corresponds well to synapse-count reductions reported for (quietly-raised) mice older than 108 weeks (Parthasarathy et al., 2018).

A number of human studies on EFRs of subcortical origin also report steady response declines with age (Goossens et al., 2016; Presacco et al., 2016; Goossens et al., 2019; Garrett & Verhulst, 2019). However, EFRs to more complex stimuli (e.g. to speech-in-noise tokens), or from more cortical generators, do not show such overt declines (Schoof & Rosen, 2016; Presacco et al., 2016), which has led several authors to conclude that encoding to temporal envelopes might be enhanced in older listeners (for an overview, see Parthasarathy et al., 2019a) due to post-SNHL changes in the midbrain (Parthasarathy et al., 2019b). The present study focused on EFRs of subcortical orgin and shows that enhanced EFRs can theoretically occur due to OHC damage. These enhancements were observed even when the stimulation level was kept fixed, and were related to the use of envelope shapes that resulted in clear ANF peak-rate and EFR magnitude increases in response to OHC deficits (e.g., SAM, H10 stimuli). A rare EFR study with recordings from young HI subjects (Goossens et al., 2019), confirms our predicted SAM-EFR enhancements for OHC damage, by showing enhanced 80-Hz ASSR SNRs for yHI compared to yNH groups for equal-level stimulation. At the same time, they observed a consistent decline in ASSR SNR for the middle-age and older groups, irrespective of whether they had NH or HI audiograms. These data, along with our own observations, and model simulations point to an interplay between the OHC and purely-age related (or synaptopathy-induced in our model simulations) aspect of SNHL which affects the EFR magnitude. Use of stimuli that are more sensitive to synaptopathy, and only minimally affected by OHC damage (e.g. the RAM stimuli) or cortical processing, can hence benefit future studies in clarifying the respective roles of cochlear synaptopathy and potential midbrain changes in temporal envelope coding along the auditory pathway.

### Implications for clinical diagnostics and sound perception studies

Finding an AEP-based metric with differential sensitivity to synaptopathy, even in the presence of OHC deficits, is an important pursuit which requires a multi-center and interdisciplinary approach. On the one hand, there is compelling evidence from animal studies that ABR wave-I amplitudes and SAM EFRs are compromised after histologically-verified synaptopathy (Kujawa & Liberman, 2009; Bourien et al., 2014; Sergeyenko et al., 2013; Shaheen et al., 2015; Möhrle et al., 2016a; Chambers et al., 2016; Parthasarathy et al., 2018), but on the other, little is known about the respective roles of OHC-damage and synaptopathy aspects in this degradation. There is a present absence of experimental approaches which vary the degree of OHC and synaptopathy damage in a controlled way. Animal studies of synaptopathy often focus on individual ABR/EFR markers after controlled synaptopathy-induction (e.g. quietly-raised animals with age and ototoxic effects) and most human studies focus on clinically normal-hearing subjects (e.g., Kujawa & Liberman, 2009; Mehraei et al., 2016; Prendergast et al., 2017a; Guest et al., 2017). Lastly, human AEP studies on listeners with impaired audiograms have the drawback that it is presently not possible to measure the true SNHL histopathology.

To bridge this translational gap, model-based approaches can have a pivotal role. Even though modeling studies are inherently limited by the quality of the adopted model, they can be effective in narrowing down the parameter space of potentially sensitive AEP markers. Promising candidate markers can afterwards be tested more efficiently in experiments with humans and animal models. Despite the theoretical starting-point we took in our stimulus design, our model predictions of how AN fibers respond differentially to stimulus type and SNHL alterations provided confidence that the population EFR response would follow these trends. Even though the model simulates the functional signal representation along the auditory pathway without modeling all specific brainstem neuron types, it has previously shown its merit at reasonably and collectively simulating the level-dependence of human OAEs, ABRs, and EFRs while accounting for the level-dependence and adaptation properties of single unit-ANF fibers (Altoé et al., 2018; Verhulst et al., 2012, 2015, 2018a; Keshishzadeh et al., 2020). It is hence not incidental that the predicted changes in EFR strength due to stimulus envelope changes, or their sensitivity to different aspects of SNHL showed a good resemblance to the trends observed experimentally.

Particularly, we demonstrated that RAM25 EFRs were more effective at identifying individual age-related SNHL differences than either the SAM EFR or the ABR waves, and this finding is important in a couple of ways. First, overall stronger EFRs improve their application range towards listeners with more severe SNHL pathologies. Whereas the SAM EFRs did not reveal group differences between the oNH and oHI listeners, the RAM EFR did, and can hence offer a more fine-grained estimate of the degree of synaptopathy in listeners with normal or impaired audiograms. Aside from a few studies that used transposed-tone EFRs with sharp envelopes (e.g., Bharadwaj et al., 2015; Guest et al., 2017), most human studies used less sensitive, smaller-amplitude markers such as the ABR wave-I amplitude or SAM-EFR to study how individual physiological responses relate to sound perception. We showed earlier that OHC damage can confound the ABR wave-I marker (Verhulst et al., 2016), and also the type of response analysis (i.e. f_1_ vs f_1_+harmonics vs noise floor correction) can introduce an inherent variability in candidate physiological markers of synaptopathy. For this reason, it would be worthwhile to re-analyse the SAM-EFRs of prior studies using the proposed analysis method to study relationships to individual sound perception, or adopt RAM stimuli in future human synaptopathy studies with listeners who may have OHC damage.

Secondly, the communality analysis showed that RAM EFRs were more sensitive to age-related aspects of SNHL and less sensitive to the co-existing OHC damage aspects. Our model simulations assign these age-related declines in EFR strength to synaptopathy. Even though the causality of this relationship should ideally be confirmed in animal studies of synaptopathy and OHC damage, it is clear that when confirmed, the RAM25 EFR might help therapeutic interventions or studies which aim to study the perceptual consequences of synaptopathy. It is known that aging causes both OHC damage (ISO-7029:2000, 1991) and synaptopathy (Sergeyenko et al., 2013; Parthasarathy et al., 2018; Wu et al., 2018), and hence older people with diagnosed OHC damage are expected to suffer from synaptopathy. This means that when audiometric thresholds predict outcomes on psychoacoustic tasks in aging studies, there is a possibility that the co-existing synaptopathy aspect of SNHL was responsible for driving the reduction in task performance.

## Conclusion

We combined model simulations with EFR recordings in normal and hearing impaired listeners to develop AEP-based stimuli which showed an enhanced sensitivity to the (age-related) synaptopathy aspect of SNHL. We conclude that supra-threshold RAM stimuli with duty cycles between 20- 25% and modulation rates around 120 Hz are maximally efficient to yield a strong response magnitude which is maximally sensitive to the synaptopathy aspect of SNHL. RAM25 magnitudes were considerably larger than commonly-used SAM-EFR markers of synaptopathy, and showed more pronounced age-related differences than ABR markers. Improving the analysis method to include the harmonics and perform a noise-floor correction further improved the sensitivity of the RAM-EFR metric. Taken together, we hope that the outcomes of this theoretical-experimental study will improve the interpretation possibilities of future studies that wish to identify the role of synaptopathy/deafferentation for sound perception as well as yield a set of sensitive markers of cochlear synaptopathy for use in animal and human studies.

## Abbreviations

ABR: auditory brainstem response
AEP: auditory evoked potential
AM: amplitude modulation
ANF: auditory-nerve fiber
ASSR: auditory steady-state response
BB: broadband
BM: basilar membrane
CF: characteristic frequency
CN: cochlear nucleus
EFR: envelope following response
H/M/LSR: high/medium/low spontaneous rate
HI: hearing-impaired
IC: inferior colliculus
IPI: inter-peak interval
IHC: inner-hair-cell
MD: modulation depth
NF: noise floor
NH: normal-hearing
OAE: otoacoustic emission
OHC: outer hair cell
peSPL: peak-equivalent sound pressure level
RAM: rectangular-wave amplitude-modulated
RMS: root mean square
SAM: sinusoidal amplitude-modulated
SNHL: sensorineural hearing loss

## Acknowledgements

This work was supported by the German Research Foundation (PP1608, VE924/1-1; VV, SV), the European Research Council (ERC-StG-678120, RobSpear; VV, SV) and DFG Cluster of Excellence (EXC 1077/1, “Hearing4all”; MG, MM). The authors would like to thank the study participants as well as the Oldenburg Hörzentrum for their help with subject recruitment.

## Author Contributions

VV: conceptualization, methodology, software, validation, formal analysis, investigation, data curation, writing: original draft, visualization; MG: methodology, software, investigation; MM: methodology, software, investigation; SV: conceptualization, methodology, resources, writing: original draft, visualisation, review & editing, supervision, project administration, funding acquisition.

## Data availability

The model code used for the simulations is available via 10.5281/zenodo.3717800 or github.com/HearingTechnology/Verhulstetal2018Model. Software to generate the stimuli and extract the EFR magnitudes from raw-EEG recordings is available on github.com/HearingTechnology/EFR2020 stimuli analysis.

